# From bats to humans: uncovering *ISG15* as a new resistance factor in type 1 diabetes

**DOI:** 10.64898/2026.04.02.713107

**Authors:** Eugenia Martin-Vazquez, Xiaoyan Yi, Maressa Fernandes Bonfim, Sayro Jawurek, Priscila L. Zimath, Arturo Roca-Rivada, Junior Garcia Oliveira, Jose Maria Costa-Junior, François Pattou, Julie Kerr-Conte, Montserrat Nacher, Eduard Montanya, Erwin Ilegems, Johnna D. Wesley, Alexandra C. Title, Burcak Yesildag, Tzachi Hagai, Anne Op de Beeck, Decio L. Eizirik

## Abstract

Viral infections are one of the main environmental factors triggering type 1 diabetes (T1D). Pancreatic alpha cells are more resistant than beta cells to diabetogenic viruses, partially explaining their survival in T1D. Similarly, bats have enhanced viral resistance, suggesting putative convergent evolution in antiviral mechanisms. Herein, we compared global gene expression in bat fibroblasts under basal conditions or exposed to double-stranded RNA to human alpha and beta cells and found that alpha cells exhibit greater similarity than beta cells to the antiviral responses of bat cells, as well as stronger intrinsic resistance to viral infection. Interferon-stimulated gene 15 (ISG15), a key regulator of antiviral responses in humans and bats, has higher expression in alpha compared to beta cells in five single-cell RNASeq datasets from human islet cells and in human induced pluripotent stem cell (hiPSC)-derived alpha-like cells. *ISG15* knockdown in human insulin-producing EndoC-βH1 cells and human islets increases apoptosis under basal conditions and after IFNα exposure, exacerbates IFNα responses and increases cell death and viral production after infection with the diabetogenic virus coxsackievirus B1, while its overexpression protects EndoC-βH1 cells from the virus. Collectively, the present results demonstrate that alpha cells but not beta cells have similarities with the virus resistance gene program present in bats and identify *ISG15* as an important factor for islet cells to cope with viral and diabetogenic stresses.

## INTRODUCTION

A surprising feature of type 1 diabetes (T1D) is that the immune system destroys insulin-producing beta cells but not glucagon-producing alpha cells in the pancreas, despite both cell types being exposed to inflammatory mediators and becoming dysfunctional (1, 2). Additionally, dysfunction often progresses to beta cell death, while alpha cells survive. We previously showed important differences between human beta and alpha cells, including divergent expression of proteins involved in the regulation of apoptosis, endoplasmic reticulum stress and the interaction between target cells and CD8+ T-cells (3–7). However, the ultimate mechanisms that explain alpha cell resistance to immune-induced beta cell damage remain to be clarified.

Viral infections are one of the potential environmental factors triggering T1D (8, 9). The enterovirus coxsackievirus (CVB) is detected in beta cells from approximately 70% of people with T1D, while present in less than 15% of non-diabetic individuals (9, 10). We previously reported that alpha and beta cells express similar amounts of the Coxsackievirus and Adenovirus Receptors (CAR) and get infected by CVB5. Antiviral mechanisms, however, are more efficient in alpha compared to beta cells, enabling alpha cells to get rid of the virus while beta cells die (5). Moreover, expression of antiviral mechanisms is higher in alpha- than in beta-like cells differentiated from human induced pluripotent stem cells (hiPSC) under basal conditions or after exposure to the pro-inflammatory cytokine IFNα (3). Notably, similar antiviral-related features appear to be upregulated in uninfected cells across multiple bat species (11–13). Bats are natural reservoirs for a wide range of viruses, e.g., coronaviruses, filoviruses, paramyxoviruses and rhabdoviruses, and unlike humans, who often experience severe inflammatory responses to viral infections, bats often exhibit asymptomatic infections (14, 15). These similarities between pancreatic alpha cells and bats suggest the presence of convergent evolutionary mechanisms that are worth exploring.

Interferon-stimulated gene 15 (ISG15) is a ubiquitin-like protein strongly induced by type I interferons (IFN-I) during viral infections in humans (reviewed in (16)) and is a key antiviral effector in bats (15). Intracellularly, free ISG15 can be conjugated to hundreds of newly synthesized host and viral proteins (via a process termed ISGylation), helping to restrict viral replication, viral protein production and viral packaging (17). Protein ISGylation is counteracted by another interferon-stimulated gene, namely ubiquitin-specific protease 18 (USP18), that is critical for negative regulation of IFN-I immunity (16). In T1D, the IFN-I signalling is key in the initial stages of insulitis and immune infiltration (18–20) but whether ISG15 and/or USP18 play a role in alpha/beta cells in T1D remains to be determined.

Herein, we compared the transcriptional gene signatures of alpha versus beta cells with the transcriptional changes induced by the double-stranded RNA poly(I:C) in bat fibroblasts to explore the similarities between human pancreatic cell populations and bat antiviral responses. We found that *ISG15,* a key regulator of bats’ antiviral responses, is upregulated in alpha compared to beta cells according to five single-cell (sc)RNASeq datasets of human islet cells and confirmed an increase of *ISG15* expression and secretion in purified alpha-versus beta-like cells differentiated from hiPSC and exposed to IFNα. We further explored the role of *ISG15* in human islet cell models, namely the pure beta cell line EndoC-βH1 (21) and native human islets under inflammatory stress conditions and viral infection. The findings obtained suggest a surprising convergent evolution in antiviral responses between bats and human alpha cells and a key role for *ISG15* in this process.

## MATERIALS AND METHODS

### Bulk RNA-seq analysis

Raw FASTQ files for FACS-sorted human alpha and beta cells were obtained from the Human Pancreas Analysis Program (HPAP) data portal (https://hpap.pmacs.upenn.edu/). Sequencing reads were subjected to quality control using fastp v0.19.6 (22), which included adapter removal, trimming of low-quality bases, and exclusion of reads shorter than 50 bp (half of the original 100 bp read length). To maintain stringent data quality and reduce confounding effects, only samples with >= 30 million high-quality reads were included. Additionally, we limited the analysis to donors with matched alpha and beta cell samples. Applying these filters, we included 12 donors in the final analysis: HPAP-006, HPAP-014, HPAP-019, HPAP-022, HPAP-026, HPAP-035, HPAP-040, HPAP-052, HPAP-053, HPAP-054, HPAP-056, and HPAP-080 (**Table S4**). Cleaned reads were aligned to the human transcriptome reference (GENCODE v36, GRCh38.p13) using Salmon v1.4.0 (23), with the options “--seqBias”, “--gcBias”, and “--validateMappings” enabled to correct for potential biases. The Salmon index was built using standard parameters. Transcript abundance estimates were imported by the tximport package (24) and summarized to gene-level counts using the summarizeToGene function. Differential expression analysis was conducted with DESeq2 v1.38.3 (25), employing the *Wald test* for statistical inference and applying Benjamini-Hochberg to adjust for multiple testing. Genes with adjusted P-value below 0.05 were considered as significantly differentially expressed.

### Rank-rank Hypergeometric Overlap (RRHO) analysis

For the global transcriptomic comparison between *Rousettus* fibroblast cells and alpha and beta cells, we employed RedRibbon analysis (26), an improved RRHO pipeline with enhanced accuracy for detecting the most significant overlapping genes. The input for this analysis consisted of two complete gene lists with log2 fold change (log2FC) calculated from differential expression analysis: bat fibroblasts exposed to poly(I:C) for 4h versus control (n=15), retrieved from the original study (27), and alpha versus beta cells under basal conditions (n=12). Briefly, log2FC values from each dataset were independently ranked along the two coordinates (i.e., x and y) using genes detected in both datasets. An evolutionary algorithm was employed to identify the gene rank indices associated with the most significant hypergeometric P-value, optimizing overlap fitness. For each quadrant, the maximal log P-values and corresponding permutation-adjusted P-values are reported. Based on this test, we determined the rank index positions with the most significant overlap and defined the most significantly co-regulated genes as those lying above the intersecting thresholds in each quadrant.

### Functional enrichment of co-regulated genes from RRHO analysis

To evaluate the functional enrichment of genes showing significant overlap in the RedRibbon comparison analysis, we performed hypergeometric overrepresentation testing using R packages ReactomePA V1.42.0 (28) and clusterProfiler V4.6.2 (29). The analysis was conducted against both the Reactome and Kyoto Encyclopedia of Genes and Genomes (KEGG) pathway reference gene sets, downloaded from MSigDB (https://data.broadinstitute.org/gsea-msigdb/msigdb/release/7.2; c2.cp.reactome.v7.2.symbols.gmt and c2.cp.kegg.v7.2.symbols.gmt versions), using the enrichPathway and enrichKEGG functions, respectively. Gene sets with a size range of 15 to 500 genes were included, and 10,000 permutations were applied to assess the robustness of the enrichment. Pathways with adjusted P-values <0.05 (Benjamini-Hochberg correction) were considered significantly enriched.

### Virus resistance and tolerance programs alignment

First, genes commonly detected in both human alpha vs beta cells and bat fibroblasts (poly(I:C) vs control) from a previous study (27) were identified. Among these intersecting genes, the top 300 upregulated and downregulated genes were defined as alpha-cell- and beta-cell-specific genes based on the differential analysis of alpha versus beta cells (**Table S2**). Second, genes primarily associated with viral tolerance and resistance programs were retrieved from a previous study (27) that integrated transcriptional responses and physiological measurements of influenza virus infection across 33 mouse strains. In this study, multidimensional data were reduced to two dimensions representing the tolerance and resistance axes, providing a relative position for all studied genes, and correlation coefficients were calculated between each gene and the two programs. Additional analyses suggest that these gene sets are conserved across mammals and infectious diseases. Third, we mapped the alpha cell- and beta cell-specific genes to extract their relative positions and coefficient values and compared them using a two-tailed Mann-Whitney test. The detailed tolerance and resistance data are provided in **Table S3**.

### Single-cell RNA-seq analysis

For the human islet datasets, raw scRNA-seq FASTQ files (10X Genomics platform) were obtained from the HPAP portal (https://hpap.pmacs.upenn.edu) (**Table S4**). The HPAP-young and HPAP-older donor groups represent control cohorts matched to T1D and T2D patients, respectively, curated by the HPAP consortium based on comparable age and body mass index (BMI) profiles. Our analysis included 15 young and non-diabetic HPAP donors (IDs: HPAP-022, HPAP-026, HPAP-027, HPAP-034, HPAP-035, HPAP-036, HPAP-037, HPAP-039, HPAP-040, HPAP-044, HPAP-047, HPAP-056, HPAP-082, HPAP-099 and HPAP-104), 13 older non-diabetic HPAP donors (IDs: HPAP-052, HPAP-053, HPAP-054, HPAP-059, HPAP-063, HPAP-074, HPAP-075, HPAP-077, HPAP-080, HPAP-093, HPAP-101, HPAP-103 and HPAP-105), 9 AAB1+ donors (HPAP-024, HPAP-029, HPAP-038, HPAP-045, HPAP-049, HPAP-050, HPAP-072, HPAP-092 and HPAP-114), 2 AAB2+ donors (HPAP-043 and HPAP-107), and 11 T1D donors (HPAP-020, HPAP-021, HPAP-023, HPAP-032, HPAP-055, HPAP-064, HPAP-071, HPAP-084, HPAP-087, HPAP-113 and HPAP-123). Clinical metadata for these donors can be accessed directly via the HPAP portal (https://hpap.pmacs.upenn.edu/explore/download?donor) using the listed IDs. Additionally, scRNA-seq FASTQ files from hiPSC-derived and hESC-derived islet-like cells were downloaded from the Gene Expression Omnibus (GEO) database under accession numbers GSE190726 (Chandra et al., hiPSC stage 6), GSE203384 (Szymczak et al., iPSC stage 7), and GSE20083 (hESC-derived cells matured *in vivo* following transplantation into humanized mouse), as detailed in **Table S4**.

Briefly, raw scRNA-seq data were processed using Cell Ranger (V6.1.2) (30) with standard settings. The cellranger count pipeline performed alignment to the GRCh38 human reference genome, assigned barcodes and unique molecular identifiers (UMIs), and generated a gene-by-cell expression matrix. To address ambient RNA contamination, we applied SoupX (V1.6.1) (31), using representative marker genes (*INS, GCG, SST, TTR, IAPP, PYY, KRT19* and *TPH1*) from initial clustering to estimate and remove background signals. The cleaned count matrices were then imported into Seurat v4.3.0 (32) for downstream filtering. Cells were retained if the met the following criteria: (1) at least 200 expressed genes per cell and each gene expressed in ≥3 cells; (2) exclusion of potential doublets, identified via scDblFinder (V1.12.0); (3) total gene counts per cell <9,000 and UMI counts <10,000; and (4) mitochondrial gene content ≤5%. This filtering resulted in high-quality single-cell datasets for subsequent analysis. We normalized gene expression using Seurat’s SCTransform, accounting for sequencing depth, and regressed out mitochondrial gene proportion, which typically reflects cell state or quality. The top 3,000 most variable genes were selected for principal component analysis (PCA). Sample integration was performed using Harmony (V1.2.0) (33), correcting for batch effects from sample identities and reagent kits, based on the first 50 principal components. The harmonized data were then visualized using Uniform Manifold Approximation and Projection (UMAP). Cell type annotation was performed with scSorter (V0.0.2) (34), using canonical marker genes, and further validated by manual inspection of marker gene expression in each cluster. For this study, we focused exclusively on alpha and beta cell populations under basal conditions, extracting them across five datasets. Differential expression analysis between alpha and beta cells was conducted using the Wilcoxon rank-sum test within each dataset, and P-values were corrected for multiple testing using Bonferroni adjustment. Genes with an adjusted P-value below 0.05 were considered differentially expressed.

### EndoC-βH1 cell line culture and transduction

The human pancreatic beta cell line EndoC-βH1 was kindly provided by R. Scharfmann (Institut Cochin, Université Paris, Paris, France) (21). EndoC-βH1 cells were cultured in Dulbecco’s modified Eagle medium with 5.6 mM glucose (Gibco, Thermo Fisher Scientific) as previously described (35). All the results shown for EndoC-βH1 cells refer to independent biological samples, i.e., different passages. *ISG15* and *USP18* gene expression was silenced using 2 different siRNAs per gene [Thermo Fisher Scientific]; siISG15 #1 (siRNA s18524): 5′-CCAUGUCGGUGUCAGAGCUtt-3′; siISG15 #2 (siRNA s195001): 5′-GCACCGUGUUCAUGAAUCUtt-3; siUSP18 #1 (Stealth RNAi 10620319): 5’-CCACUGGCAGGAAACUGCAUAUCUU-3’ and siUSP18 #2 (siRNA s22260): 5’-UUUUUGAUGUGGACUCAAAtt-3’. AllStars Negative Control siRNA (siCTRL) (Qiagen) was used as a negative control; the siRNA control does not interfere with beta-cell gene expression, function or viability (36). Cells were transfected using the Lipofectamine RNAiMAX lipid reagent (Invitrogen, Life Technologies) in Opti-MEM (Gibco, Thermo Fisher Scientific) and with 30 nM of each siRNA, overnight as previously described (3). After transfection, cells were kept in culture for a 24h recovery period and subsequently exposed or not to human IFNα (2,000 U/mL; PeproTech) for 24h or 48h or to viral infection (See *Viral infection and titration* section). For overexpression experiments, EndoC-βH1 cells were transduced with adeno-associated virus (AAVs) encoding *ISG15* under the CMV promoter (construct AAV/KP1-CMV-hISG15, Vector Biolabs) in standard culture media. After transduction, cells were kept in culture for a 72h recovery period and subsequently exposed or not to viral infection (*See Viral infection and titration section*).

### hiPSC-derived islet-like cells culture and FACS purification

The hiPSC line 1.023A (kindly provided by Dieter M Egli, University of Columbia) was derived from bone marrow, had a normal karyotype (XY), classical colony morphology and expressed pluripotency markers, as shown in (37–39). Cells were maintained in E8 medium (Life Technologies) in Matrigel-coated plates and were differentiated into pancreatic islet-like cells using a previously described seven-step protocol (3, 37–40). Islet-like aggregates were further differentiated by extra 4 weeks of culture (41). Enrichment of alpha-/beta-cells was done by fluorescence-activated cell sorting (FACS) using the antibodies CD26-PE for alpha-cells and CD49a-FITC for beta-cells (42, 43). Viable cells were selected using a viability dye (Fixable Viability Dye eFluor™ 780, Ebioscience). At least 50K alpha-/beta cells were collected and immunofluorescence analyses were performed to determine the enrichment and purity of the populations. Briefly, cells were fixed for 15 min in 4% PFA, washed twice with PBS, permeabilized with Triton X-100, 0.3% in PBS and blocked with UltraCruz Blocking Reagent (Santa Cruz Biotechnology). Cells were then incubated overnight with primary antibody: FLEX Polycolonal Guinea Pig Anti-Insulin Ready-to-Use (Link) (Agilent Technologies), Monoclonal Anti-Glucagon antibody produced in mouse (Sigma) and Anti-Somatostatin antibody (ab108456) (abcam) diluted in PBS. Cells were washed with PBS and incubated with secondary antibodies from Jackson ImmunoResearch: Alexa Fluor® 488 AffiniPure® F(ab’)₂ Fragment Donkey Anti-Guinea Pig IgG (H+L); Alexa Fluor® 647 AffiniPure® Donkey Anti-Mouse IgG (H+L); Alexa Fluor® 568 AffiniPure® Donkey Anti-Rabbit IgG (H+L) in PBS. Samples were mounted with Vectashield with DAPI (Vector Laboratories, Newark, CA, USA) and covered with glass coverslips. Images were acquired by widefield fluorescence microscopy (Zeiss, Oberkochen, Germany).

The rest of cells collected post-sorting were kept in culture and subsequently exposed or not to human IFNα (2,000 U/mL; PeproTech) for 24h. Then, they were processed for quantitative real-time PCR and the supernatant was collected for ELISA analyses.

### Native human islets culture and transduction

Human islets were isolated from non-diabetic organ donors by collagenase digestion and density gradient purification with the approval of the local ethical committees in Lille, France and Barcelona, Spain. The islets were characterized as previously reported (44) and cultured in M199 culture medium (5.5 mM glucose) ahead of transport to the Brussels laboratory for additional experiments. On arrival, the islets were dispersed and cultured in Ham’s F-10 medium containing 6.1 mM glucose (Gibco, Thermo Fisher Scientific), 10% fetal bovine serum (Gibco, Thermo Fisher Scientific), 2 mM GlutaMAX (Gibco, Thermo Fisher Scientific), 50 mM 3-isobutyl-1-methylxanthine (Sigma-Aldrich), 1% fatty acid–free bovine serum albumin (BSA) fraction V (Roche), and penicillin (50 U/mL) and streptomycin (50 mg/mL) (Lonza, Verviers, Belgium) (35). Immunofluorescence analyses were performed on dispersed cells to estimate the purity. Briefly, cells were fixed for 15 min in 4% PFA, washed twice with PBS, permeabilized with Triton X-100, 0.3% in PBS and blocked with UltraCruz Blocking Reagent (Santa Cruz Biotechnology). Cells were then incubated overnight with the primary antibody FLEX Polycolonal Guinea Pig Anti-Insulin Ready-to-Use (Link) (Agilent Technologies). Cells were washed with PBS and incubated with the secondary antibody from Jackson ImmunoResearch: Alexa Fluor® 488 AffiniPure® F(ab’)₂ Fragment Donkey Anti-Guinea Pig IgG (H+L) in PBS. Samples were mounted with Vectashield with DAPI (Vector Laboratories, Newark, CA, USA) and covered with glass coverslips. Insulin positive cells were manually quantified by widefield fluorescence microscopy (Zeiss, Oberkochen, Germany). All the experimental replicates are from different donors. The characteristics of the different donors are listed in **Table S5.** *ISG15* and *USP18* gene expression was silenced using the siISG15 #1 (siRNA s18524) and siUSP18 #1 (Stealth RNAi 10620319). AllStars Negative Control siRNA (siCTRL) (Qiagen) was used as a negative control. Cells were transfected using the same approach as for EndoC-βH1 cells. After transfection, cells were kept in culture for a 24h recovery period and subsequently exposed or not to human IFNα (2,000 U/mL; PeproTech) for 48h or to viral infection (See *Viral infection and titration* section).

### Human islet microtissues (hIsMTs) culture and transduction

Human islet microtissues (hIsMTs) (InSphero, MT-04-002-01-60) were generated after dispersion and reaggregation of primary human islets from a single donor (UNOS ID #ALIR128; male; 41y.o.; Hispanic; BMI 27.77; positive serologies: CMV IgG, EBV IgG; HbA1c: 5.3%; cause of death: head trauma). hIsMTs were transduced with AAVs encoding specific shRNA targeting *ISG15* (siISG15 #1 sequence) from Vector Biolabs during aggregation for 5 days in Akura hanging drop plates (InSphero, CS-PF24) and then released into Akura 96-well plates (InSphero, CS-09-001-00). hIsMTs were cultured in standard culture medium (InSphero, CS-07-005-01) or standard culture medium containing cytokines: TNFα (25 ng/mL, R&D Systems, 10291-TA-050); IL1β (5 ng/mL, Sigma, H6291-10UG); IFNγ (25 ng/mL, R&D Systems, 10067-IF-100) and IFNα (10 ng/mL, Peprotech, 300-02AA) starting at day 9 after reaggregation, for 6 days. Medium was exchanged and cytokines were re-dosed every 2-3 days throughout the experiment. This combination and concentration of cytokines together with the time of exposure were selected in previous dose-response and time-course experiments to induce severe dysfunction and apoptosis in hIsMTs (45–47).

### Cell viability assessments

EndoC-βH1 cells and human islets viability was determined by fluorescence microscopy using the nuclear dyes propidium iodide (5 μg/mL; Sigma-Aldrich) and Hoechst 33342 (10 μg/mL; Sigma-Aldrich), as previously described (36, 48). Viability was evaluated by two independent observers, one of them being unaware of sample identity. The agreement between the two observers was >90%. The results are expressed as percentage of apoptosis, calculated as the number of apoptotic cells/total number of cells and presented as fold change compared to siCTRL cytokines-treated cells.

### Quantitative real-time PCR

EndoC-βH1 cells, hiPSC-derived islet-like cells and human islets were washed with PBS, detached and collected in lysis buffer. Poly(A) + RNA was isolated using the Dynabeads mRNA DIRECT Kit (Invitrogen) and reverse-transcribed using the Reverse Transcriptase Core Kit (Eurogentec), following the manufacturer’s protocol. hIsMTs were pooled (6 per well), washed twice in PBS and transferred to PCR-clean 2 mL Eppendorf tubes. QIAzol lysis buffer (Qiagen, 79306) was added and hIsMTs were vortexed and sonicated to aid thorough lysis. The lysate was snap-frozen in liquid nitrogen and total RNA was extracted using the RNeasy Micro Kit (Qiagen) and reverse-transcribed using the Reverse Transcriptase Core Kit (Eurogentec). The quantitative RT-PCR amplification reactions were performed with IQ SYBR Green Supermix (Bio-Rad Laboratories) and ran in the CFX Connect Real-Time PCR Detection System (Bio-Rad Laboratories). The product quantification was performed using the standard curve method (49) or the 2(-Delta Delta C(T)) Method (50). For each gene, the melting curve was analysed to confirm amplification of a single PCR product. Gene expression values were normalized by the geometric mean of the housekeeping genes *ACTIN* and *VAPA* (51) and presented as fold change as detailed in the figure legends. The primers used are listed below: *ACTIN*; FW: CTGTACGCCAACACAGTGCT, RV: GCTCAGGAGGAGCAATGATC, *ARX*; FW: CTGCTGAAACGCAAACAGAGGC, RV: CTCGGTCAAGTCCAGCCTCATG, *BIM*; FW: TTCTTGCAGCCACCCTGC, RV: CTTGCGTTTCTCTCAGTCCGA, *CXCL10*; FW: GTGGCATTCAAGGAGTACCTC, RV: GCCTTCGATTCTGGATTCAG, *DP5;* FW: GAGCCCAGACTTGAAAGG, RV: CCCAGTCCCATTCTGTGTTT, *HLA-ABC;* FW: CAGGAGACACGGAATGTGAA, RV: TTATCTGGATGGTGTGAGAACC, *ISG15;* FW: ACAAATGCGACGAACCTCTG, RV: CGCAGATTCATGAACACGGT*, PDX1;* FW: AAAGCTCACGCGTGGAAA, RV: GCCGTGAGATGTACTTGTTGA, *PUMA;* FW: TTGTGCTGGTGCCCGTTCCA, RV: AGGCTAGTGGTCACGTTTGGCT, *USP18;* FW: CAATCCACCTCATGCGATTCT, RV: TTGGAAGGATCTGGCTGAAATC, *VAPA*; FW: TACCGAAACAAGGAAACTAATGGAA, RV: GCCTTAAACCTTCATCTCTCAGGT.

### Protein extraction and Western blot analysis

Total protein from EndoC-βH1 cells was extracted using Laemmli buffer supplemented with phosphatase and protease inhibitors (Roche) and separated on 10% SDS–PAGE. The nitrocellulose membranes were probed using specific primary antibodies: ISG15 Polyclonal antibody Rabbit #15981-1-AP Proteintech, USP18 (D4E7) Rabbit mAb #4813 Cell Signaling, Human GAPDH Antibody (Bio-Techne Sales Corp. 2275-PC-100) or Anti-TUBA4A (TUBA1) Antibody #T5168 Sigma diluted 1:1,000 in TBST (TBS, 0.1% Tween 20) with 5% BSA. The densitometric values were quantified by ImageLab software version 6.1 (Bio-Rad Laboratories, RRID:SCR_014210) and normalized to TUBULIN or GAPDH after background subtraction.

### CXCL10 and ISG15 secretion by ELISA

The cell culture supernatants were collected at the end of the experiments and immediately stored at −80°C until sample processing. The assay procedures and calculation of the results were conducted following the manufacturer’s recommendation. Human CXCL10/IP-10 Immunoassay (Quantikine ELISA kit, R&D Systems) was used to quantify CXCL10 secretion to the culture medium as indicated in the figure legends. Human ISG15 ELISA Kit (Abcam) was used to quantify ISG15 secretion to the culture medium as indicated in the figure legends.

### Viral infection and titration

The prototype strain of enterovirus, CVB1/Conn-5, was acquired from the American Type Culture Collection (ATCC, Old Town Manassas, VA, United States). This virus was propagated in green monkey kidney (GMK) cells. EndoC-βH1 cells and human islets were infected with CVB1 at indicated multiplicity of infection (MOI) based on previous publications (5, 52) in EndoC-βH1 or human islets media without BSA and containing 2% FBS. After a 2h adsorption period at 37°C, the inoculum was removed. Culture medium was then added, allowing the virus to replicate for the specified durations. Viral production in the medium of infected cells was assessed by titration using endpoint dilutions in microwell cultures of GMK cells. After 24h, cytopathic effects were observed via microscopy as described below, and 50% tissue infection dose titers were calculated using the Kärber formula (53). Relative viral titers were expressed as the ratio between viral titers in the siCTRL/NT and siISG15/AAV-ISG15 conditions.

### Assessment of cytopathogenic effect

The percentage of viable and apoptotic cells was determined after staining with the above-mentioned DNA-binding dyes Hoechst 33342 (10 μg/mL) and propidium iodide (5 μg/mL). Viability was evaluated by two independent observers, one of them being unaware of sample identity. The agreement between the two observers was >90%. Results are expressed as the percentage of apoptosis.

### 3D staining and imaging

hIsMTs were washed twice in PBS (with Mg ^+^Ca ^+^), fixed for 15 min in 4% PFA, washed twice more with PBS and kept in PBS with 0.05% sodium azide until staining. hIsMTs were then permeabilized with permeabilization buffer (Triton® X-100, 0.5% in PBS w/o Mg ^+^Ca ^+^) and washed twice with PBS. hIsMTs were blocked with 5% donkey serum in PBS to prevent nonspecific antibody binding. hIsMTs were then incubated overnight with primary antibody: rabbit anti-NKX6.1 (Abcam, ab221549) and sheep anti-ARX (R&D Systems, AF7068-SP) in antibody dilution buffer (5% donkey serum, 0.2% Triton® X-100 in PBS w/o Mg ^+^Ca ^+^). hIsMTs were washed with wash buffer (0.2% Triton® X-100 in PBS w/o Mg2^+^Ca2^+^) and incubated with secondary antibodies (donkey anti-rabbit AF568 (Thermofisher, A10042); donkey anti-sheep AF647 (Jackson ImmunoResearch, 713-605-147) in antibody dilution buffer, along with DAPI (D9542). hIsMTs were washed 3 times with wash buffer, transferred into Akura 384-well ImagePro plates (InSphero, CS-PC14), and cleared with ScaleS4 solution for 3D imaging. Images were acquired using the Yokogawa CQ1 Benchtop High-Content Analysis System, taking fluorescent images in 3 µm z-steps. Images were quantified using customized CellPathfinder pipelines.

### Replicates

In each experiment, n=1 corresponds to one independent biological observation, i.e. EndoC-βH1 cells from different passages, hiPSC-derived islet-like cells from different differentiations or human islets from different donors. The experiments performed in hIsMTs correspond to a single donor and are considered as indicative; the technical replicates are shown.

### Statistics

Data are presented as mean ± Standard Error of the Mean (SEM) and analyzed by unpaired Student’s t-test or one-way ANOVA followed by Bonferroni post-hoc correction for multiple comparisons, as indicated in the figure legends. P-values less than or equal to 0.05 were considered statistically significant. Grubbs method was used to identify data outliers. Sample size is displayed in each graph and it was calculated based on the variability observed in preliminary experiments and power analysis using GPower 3.1 software. Statistical analyses were performed using GraphPad Prism software version 8 (GraphPad Software, La Jolla, USA).

## RESULTS

### Alpha cells show closer transcriptomic similarity than beta cells to the antiviral responses of bat skin fibroblasts

We compared the transcriptomes of human alpha and beta cells from 12 donors retrieved from the Human Pancreas Analysis Program (HPAP; https://hpap.pmacs.upenn.edu/) versus those of fibroblast skin cells from bats (27) by applying the Rank-Rank Hypergeometric Overlap (RRHO) analysis using the RedRibbon tool (26). The transcriptome of alpha cells showed greater similarity to bat skin fibroblasts stimulated with poly(I:C) than beta cells (**Figure 1A**). Functional enrichment analysis revealed that many of the genes commonly upregulated (457 genes; **Table S1**) in both alpha cells and poly(I:C)-stimulated bat fibroblasts were associated with interferon signaling and antiviral responses (**Figure 1B-C**), including “Interferon alpha/beta signaling”, “RIP-mediated NF-kB activation via ZBP1”, “ZBP1 (DAI) mediated induction of type I IFNs”, and “Herpes simplex virus 1 infection”. These results suggest that alpha cells (contrary to beta cells) exhibit a basal induction of antiviral-related genes.

**Figure 1.**
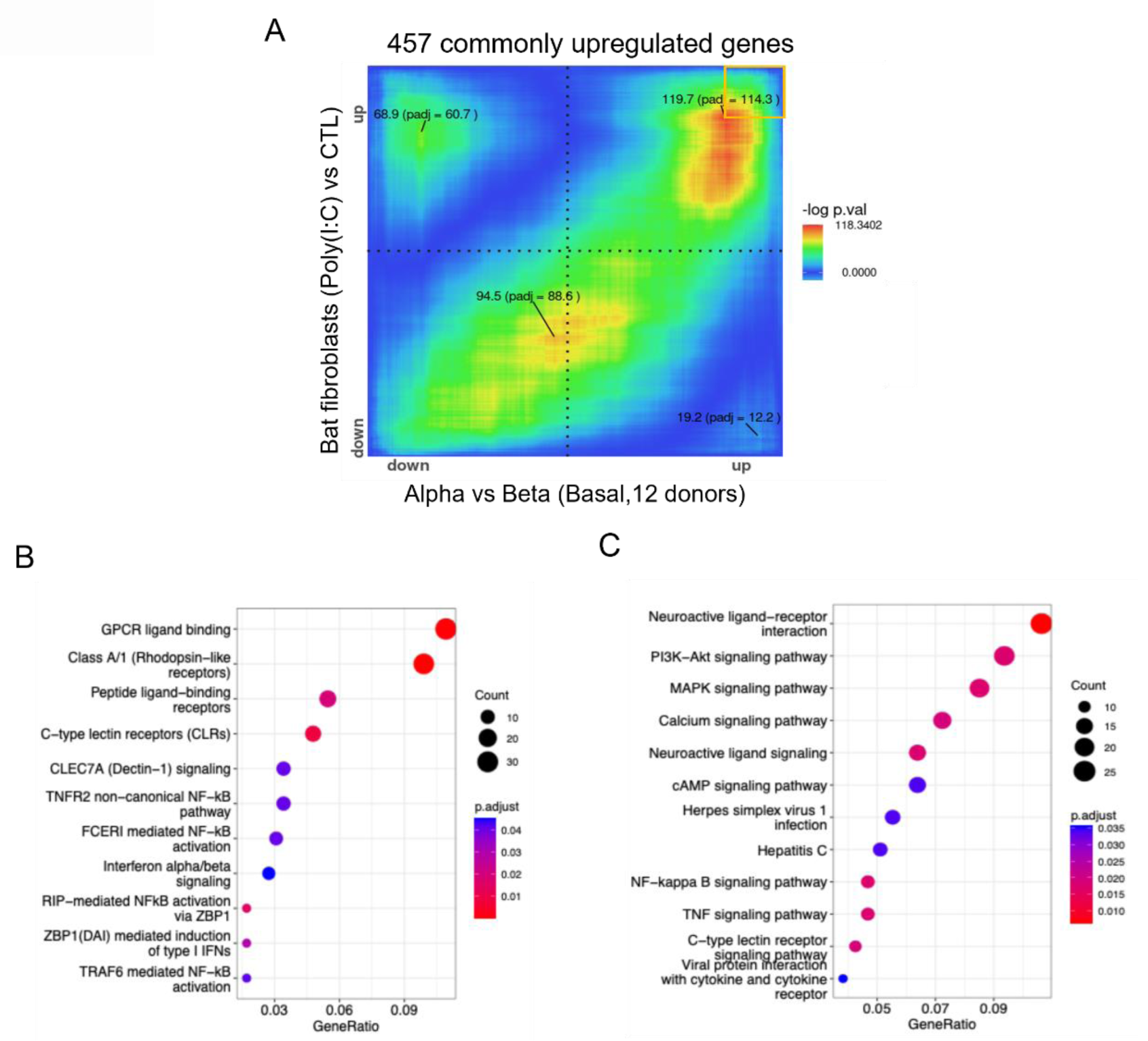
Alpha cells show closer transcriptomic similarity than beta cells to the antiviral responses of bat skin fibroblasts. **(A)** Rank-rank Hypergeometric Overlap (RRHO)-RedRibbon analysis comparing gene expression profiles between *Rousettus* skin fibroblasts treated with poly(I:C) and human alpha versus beta cells. The orange box highlights the coordinate range used to define the 457 commonly upregulated genes, based on their consistent upregulation in both comparisons. **(B)** Functional enrichment analysis of the shared upregulated genes (457 genes) from **(A)** using Reactome pathway databases. **(C)** Functional enrichment analysis of the shared upregulated genes from **(A)** using KEGG pathway databases. Gene expression data for Rousettus aegyptiacus skin fibroblasts (N=15) stimulated with poly(I:C) for 4h were obtained from the original study (27). Basal alpha and beta cell RNA-seq FASTQ files were retrieved from the Human Pancreas Analysis Program, including 12 donors with matched alpha and beta cell profiles (https://hpap.pmacs.upenn.edu/) and reanalyzed. One-to-one orthologous genes between species were used to ensure cross-species comparability.

### Alpha-cell-specific genes exhibit stronger alignment with virus resistance programs compared to beta-cell-specific genes

To investigate whether alpha- and beta-cell-specific genes (**Table S2**) are differentially associated with virus tolerance or resistance programs, we examined their distribution along the tolerance and resistance axes and their gene-to-program correlation scores as previously defined (54) (**Figure 2; Table S3)**. No significant differences were observed between alpha- and beta-cell-specific genes in either their coordinates on the tolerance axis (**Figure 2A**) or their correlation with the tolerance program (**Figure 2C**). On the other hand, alpha-cell-specific genes displayed significantly higher coordinates on the resistance axis compared to beta-cell-specific genes (P = 0.02; **Figure 2B**) and exhibited significantly stronger positive correlation with the resistance program (P = 2.2 × 10⁻⁴; **Figure 2D**). These results confirm that alpha cells, as bat fibroblasts, have features associated with antiviral resistance.

**Figure 2.**
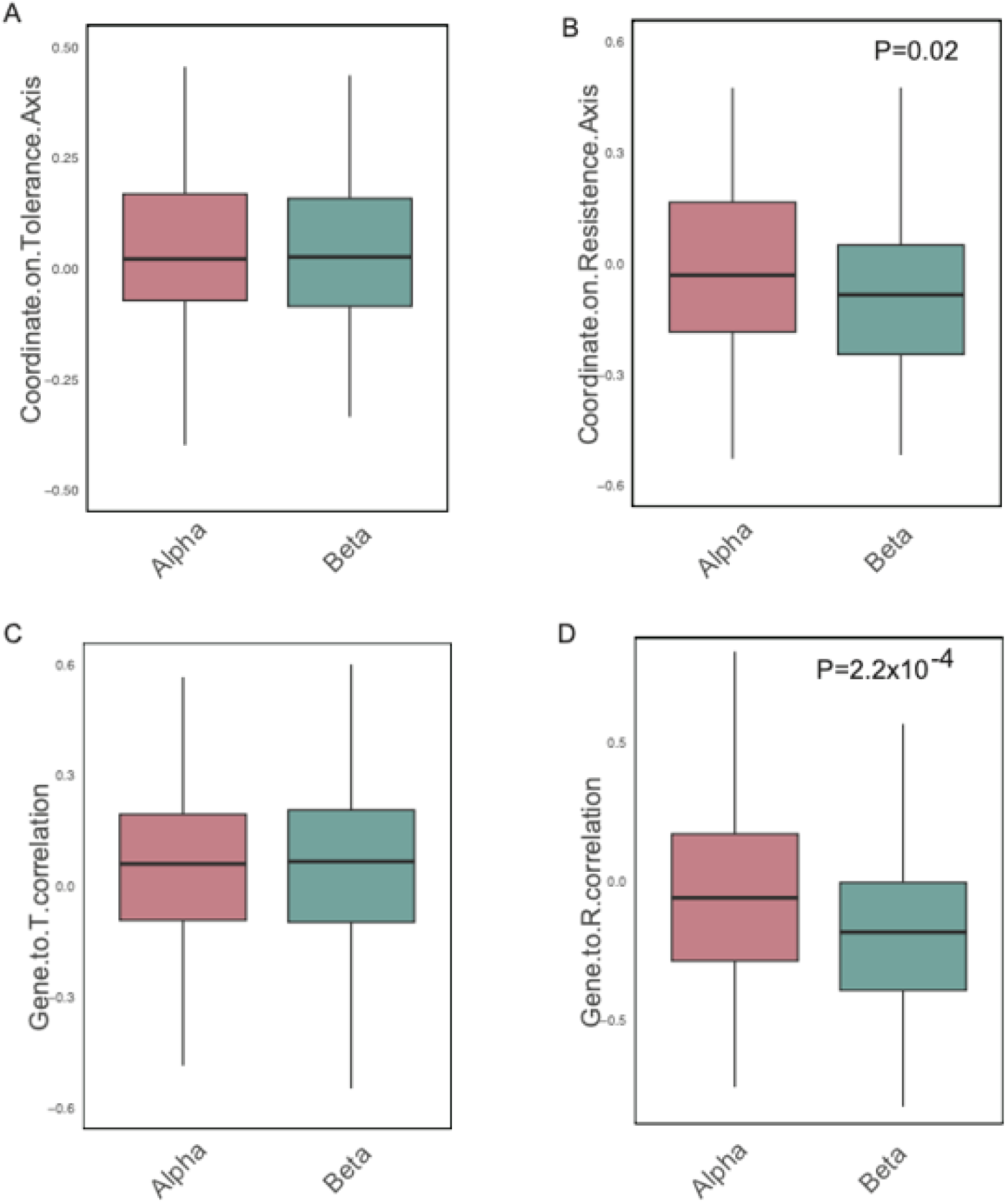
Alpha-cell-specific genes exhibit stronger alignment with virus resistance programs compared to beta-cell specific genes. (A-B) Coordinate distributions of tolerance **(A)** and resistance **(B)** axes compared between alpha- and beta cell-specific genes. **(C-D)** Correlation scores to tolerance **(C)** and resistance **(D)** programs compared between alpha- and beta cell-specific genes. Tolerance and resistance coordination scores, as well as gene-to-program correlation scores, were obtained from ^24^. Alpha- and beta-cell-specific genes were defined based on differential gene expression analysis of FACS-sorted alpha and beta cells from HPAP bulk RNA-seq data, selecting the top 300 genes upregulated in alpha cells (alpha-cell specific genes) and the bottom 300 in beta cells (beta-cell specific genes). Statistical comparisons were performed using two-tailed Mann-Whitney tests. FACS: Fluorescence-Activated Cell Sorting. HPAP: Human Pancreas Analysis Program.

### *ISG15* has higher expression in alpha compared to beta cells and its silencing in human islet cells increases apoptosis

Given the potential role of ISG15 in bats’ viral resistance (15), we next compared *ISG15* mRNA expression levels between alpha and beta cells. We re-analyzed five independent single-cell (sc)RNA-seq datasets from primary human islets from non-diabetic individuals from the HPAP (15 young, non-diabetic donors assigned as ‘HPAP-young’ and 13 older, non-diabetic donors designated as ‘HPAP-older’). HPAP-young and HPAP-older donors represent matched T1D and type 2 diabetes (T2D) controls curated by the HPAP consortium (55) (described in the Materials and Methods section). We previously showed that age has a modest impact on the transcriptional differences between alpha and beta cells (47). Additionally, we re-analyzed two hiPSC-derived islet-like cell datasets: one from cells at stage 6 of differentiation, exhibiting an immature phenotype (identified as “Chandra et al., stage 6”) (56) and the other from cells at stage 7, which is the last stage of the hiPSC-to-islet-like cells differentiation protocol (3, 37–40) (identified as “Szymczak et al., stage 7”) (3). Finally, we analyzed one human embryonic stem cell (hESC)-derived islet-like cell dataset from *in vivo* matured cells after transplantation into humanized mouse models (“Sintov et al., *in vivo*”) (57) (**Table S4**). *ISG15* expression was consistently higher in alpha cells compared to beta cells (**Figure 3A-E**). Moreover, the proportion of *ISG15*-expressing cells was significantly greater in alpha cells than in beta cells (**Figure 3F**). *ISG15* also shows higher expression in alpha cells than in beta cells from autoantibody positive (AAB+) donors (**Figure S1A, B**), but beta cells from T1D donors displayed higher *ISG15* expression as compared to alpha cells (**Figure S1C**). These results suggest that the higher basal *ISG15* expression in alpha versus beta cells reflects stronger intrinsic antiviral readiness, while in T1D cytokine-driven hyperactivation during the autoimmune attack probably leads to higher *ISG15* expression in beta cells. To confirm these findings *in vitro*, we isolated hiPSC-derived alpha-like and beta-like cells using fluorescence activated-cell sorting (FACS) (**Figure S2A**). Purity of the post-sorting cell populations was >75% by immunofluorescence staining (**Figure S2B, C**), a finding confirmed by higher *PDX1* and *ARX* expression (beta- and alpha-specific cell markers) in respectively beta- and alpha-like cells (**Figure S2D, E**). *ISG15* expression and secretion was higher in alpha-like cells vs beta-like cells as shown by qPCR (**Figure S2F**) and ELISA (**Figure S2G**) in untreated conditions and after IFNα treatment.

**Figure 3.**
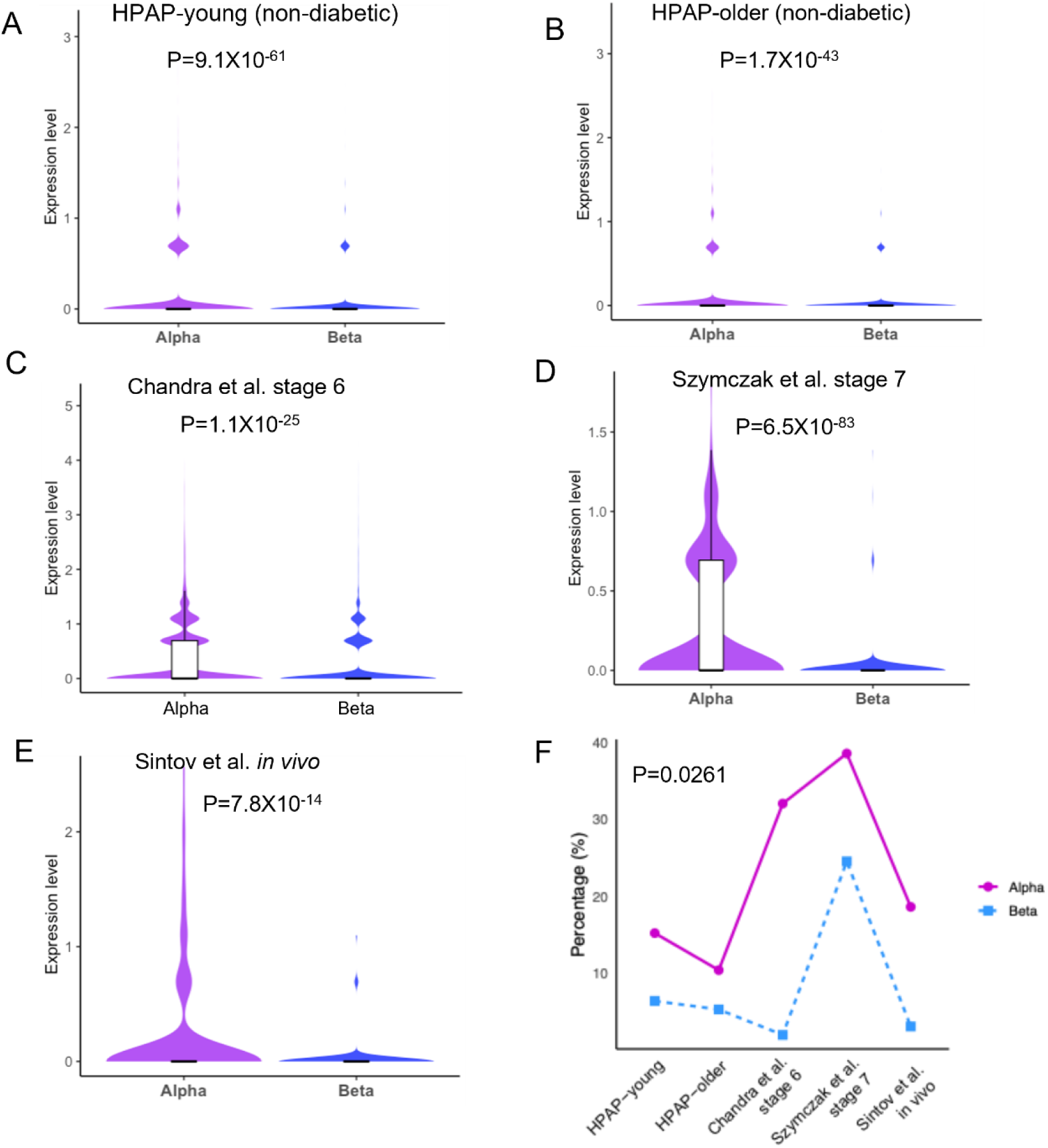
*ISG15* has higher expression in alpha cells compared to beta cells. Violin plots with an embedded boxplot from five single-cell RNA-seq datasets, including primary islets from HPAP non-diabetic donors (**A**, young; **B**, older) ^25^, hiPSC-derived endocrine cells (**C**, Chandra et al., stage 6 ^27^; **D**, Szymczak et al., stage 7 ^3^), and hESC-derived islet-like cells (**E**, Sintov et al., *in vivo*) ^32^. **(F)** Percentage of cells expressing *ISG15* by individual dataset. Plots display the expression of *ISG15* for alpha and beta cells, confirming distinct expression between both cell types in each dataset. Adjusted P-values (Wilcoxon rank-sum test with Benjamini-Hochberg correction) for panels **(A-E)** and the P-value from the paired t-test for the percentage of *ISG15*-expressing cells (panel **F**) are indicated. HPAP: Human Pancreas Analysis Program, T1D: Type 1 diabetes, T2D: Type 2 diabetes, hiPSC: human induced pluripotent stem cells, hESC: human embryonic stem cells.

We next explored the role of *ISG15* in human EndoC-βH1 cells and native human islets exposed to IFNα. Decreasing *ISG15* expression using 2 independent small interfering (si)RNAs (siISG15 #1 led to a 50-60% reduction and siISG15 #2 to a 40-50% reduction, **Figure S3A, B**) increased expression of the chemokine *CXCL10* in IFNα-exposed EndoC-βH1 cells (**Figure S3C**). For subsequent experiments, we selected the siISG15 #1 and focused the analysis on the effects of long-term exposure to IFNα, evaluated after 48h exposure to the cytokine. Silencing *ISG15* increased apoptosis under untreated conditions and after 48h of IFNα exposure in EndoC-βH1 (**Figure 4A, B**) and primary human islets (**Figure 4C, D**). Of note, in untreated conditions the expression of *ISG15* was extremely low and not significantly decreased by the silencing (see magnification of **Figure 4A, C**), suggesting that this gene is important for beta-/islet-cell integrity even in basal conditions. Additional experiments in human islet microtissues (hIsMTs) showed that *ISG15* mRNA expression was ∼500X-induced by cytokine mixtures that included IFNα versus the 100X fold induction after a treatment with IFNγ+IL1β+TNFα but without IFNα (**Figure S4A**), supporting a key role for IFNα on *ISG15* induction. Based on this, we focused on an IFNα-containing mix in subsequent experiments, where we knocked down *ISG15* using a short hairpin RNA (shRNA, **Figure S4B**). Quantitative 3D microscopy revealed a significant loss of alpha and beta cells after cytokine exposure, which was further exacerbated by *ISG15* knockdown (KD) (**Figure S4C-E**). These results confirm that *ISG15* is important for islet integrity and survival under basal and inflammatory conditions.

**Figure 4.**
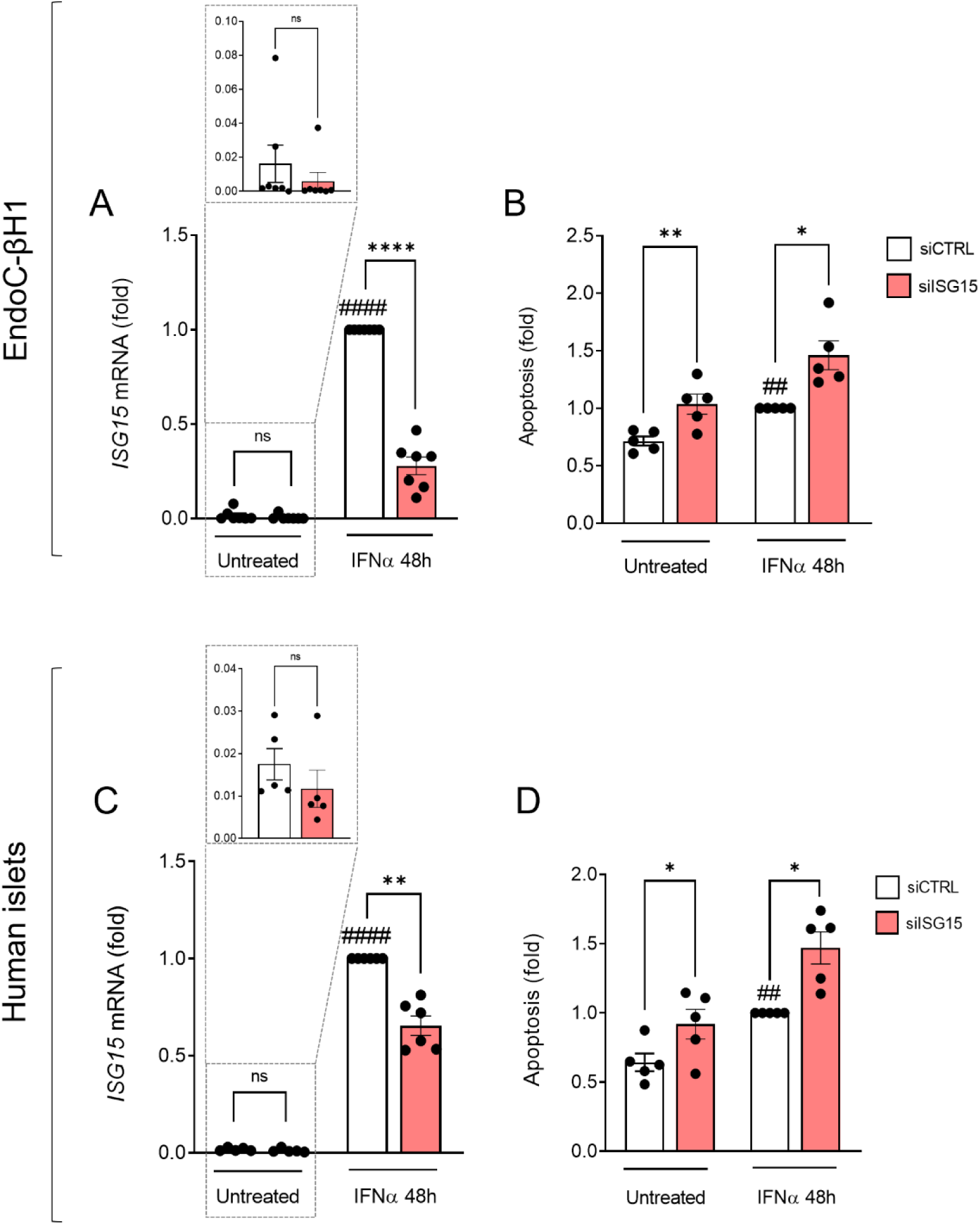
Silencing *ISG15* in human islet cells increases apoptosis. (A, **B)** EndoC-βH1 cells and **(C, D)** human islets were transfected with small interfering RNAs (siCTRL or siISG15). After recovery, cells were left untreated or exposed to IFNα (2,000U/mL) for 48h. **(A, C)** *ISG15* mRNA expression was assessed by RT-qPCR, normalized to the geometric mean of *ACTIN* and *VAPA* and presented as fold change compared to siCTRL IFNα-treated cells. **(B, D)** Apoptotic cells were counted with the DNA-binding dyes propidium iodide and Hoechst 33342. Results are means ± SEM. Each point represents an independent experiment, i.e., independent passage or islet donor. *p<0.05, **p<0.01, ****p<0.0001 vs siCTRL, ##p<0.01, ####p<0.0001 vs untreated one-way ANOVA followed by Bonferroni’s post hoc test.

### Silencing *ISG15* in human islet cells increases IFNα-induced responses

Expression and secretion of the pro-inflammatory chemokine CXCL10 is a hallmark of ‘insulitic’ islet cells and is involved in inflammation, attraction of CD8^+^ T cells and beta cell destruction (58). CXCL10 expression and secretion was increased after *ISG15* KD in IFNα-exposed EndoC-βH1 cells (**Figure 5A, B**) and human islets (**Figure 5E, F**). HLA-ABC hyperexpression at the cell surface is a hallmark of T1D in beta cells (59). Silencing *ISG15* increased IFNα-induced *HLA-ABC* expression in EndoC-βH1 cells (**Figure 5C**) and human islets (**Figure 5G**). Of note, *ISG15* KD increased *USP18* mRNA expression in IFNα-treated cells in the 2 islet-cell models studied (**Figure 5D, H**). Both *ISG15* and *USP18* are essential to ensure negative regulation of IFNα responses. Thus, we next explored the expression and role of *USP18* upon IFNα exposure. Downregulation of *USP18* (validated with 2 independent siRNAs, both leading to a 45-50% reduction and with siUSP18 #1 selected for subsequent studies, **Figure S5**) increased apoptosis in IFNα-treated EndoC-βH1 (**Figure S6A**) but induced only a trend towards increased apoptosis in human islets (**Figure S6B, C**). Notably, in human islets the KD of *USP18* mRNA was limited (25% reduction upon IFNα exposure; **Figure S6B**), potentially explaining the lack of effects on apoptosis in this model. In spite of this, IFNα-induced downstream responses were upregulated in *USP18*-silenced cells: CXCL10 expression and secretion increased significantly in EndoC-βH1 (**Figure S6D, E**) and human islets (**Figure S6H, I**). Furthermore, *USP18* KD led to increased *HLA-ABC* expression in the 2 models (**Figure S6F, J**) and upregulation of *ISG15* expression (**Figure S6G, K**). These results confirm that both genes provide important negative feedback on the IFNα responses. Previous studies showed that *USP18* inhibition in rat pancreatic beta cells exacerbates the STAT signalling pathway and increases IFN-induced apoptosis (60). Silencing *USP18* increased the expression of the IFNα-induced pro-apoptotic genes *BIM, PUMA* and *DP5* in EndoC-βH1 (**Figure S7A-C**), indicating that *USP18* plays a key role in the modulation of the intrinsic pathway of apoptosis. Notably, the KD of either *ISG15* or *USP18* increased the expression of the other at the mRNA level. At the protein level, however, silencing of *ISG15* decreased USP18 protein expression in EndoC-βH1 cells, while *USP18* silencing did not affect ISG15 protein expression (**Figure 6**).

**Figure 5.**
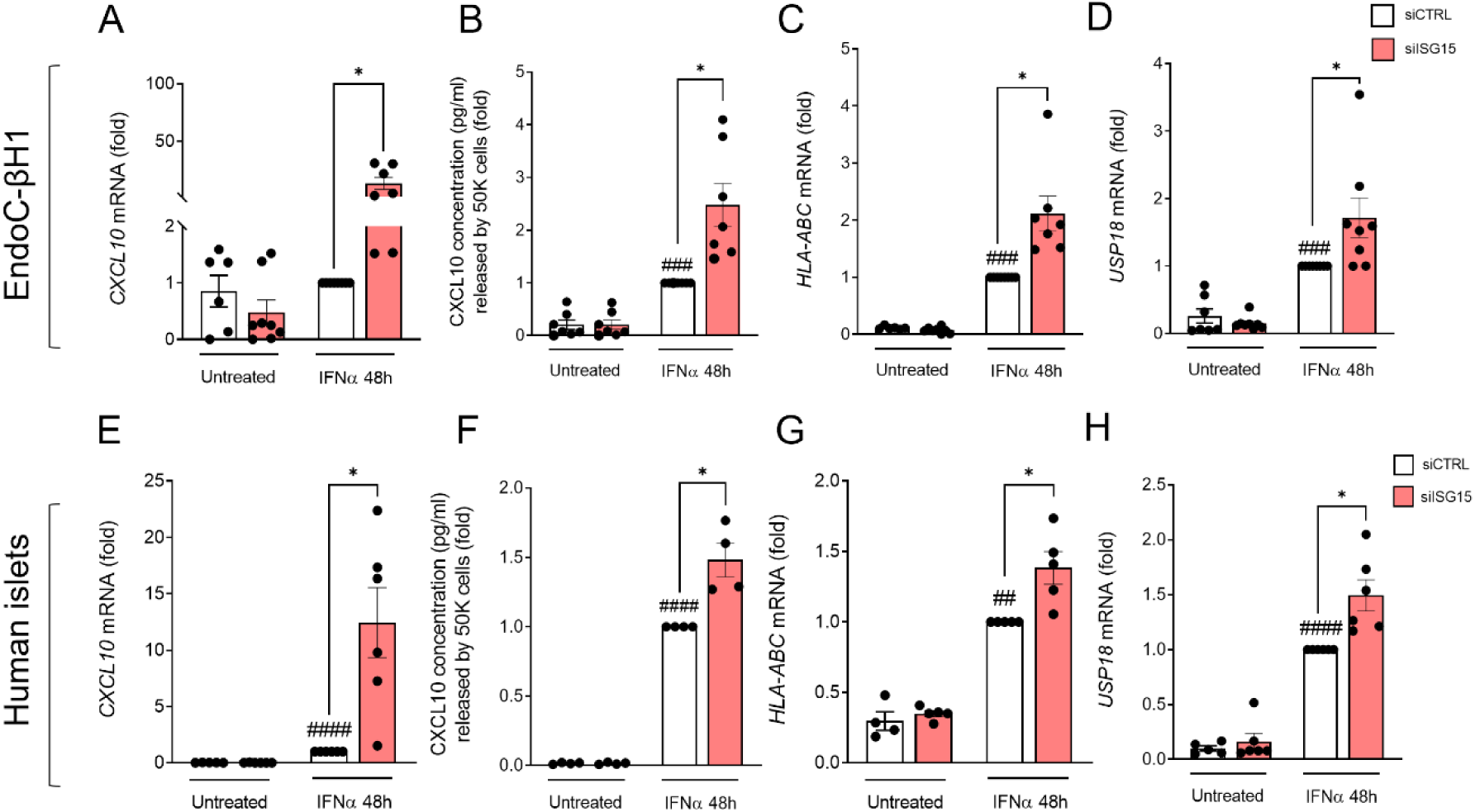
Silencing *ISG15* upregulates IFNα-induced downstream events. **(A-D)** EndoC-βH1 cells and **(E-H)** human islets were transfected with small interfering RNAs (siCTRL or siISG15). After recovery, cells were left untreated or exposed for 48h to IFNα (2,000U/mL). **(A, E)** *CXCL10***, (C, G)** *HLA-ABC* and **(D, H)** *USP18* mRNA expression was assessed by RT-qPCR, normalized to the expression of *ACTIN* and *VAPA* and represented as fold change compared to siCTRL IFNα-treated cells. **(B, F)** CXCL10 secretion in the supernatant was measured by ELISA. 50,000 cells were used per condition. Results are means ± SEM. Each point represents an independent experiment, i.e., independent passage or islet donor. *p<0.05 vs siCTRL, ##p<0.01, ###p<0.001, ####p<0.0001 vs untreated, one-way ANOVA followed by Bonferroni’s post hoc test.

**Figure 6.**
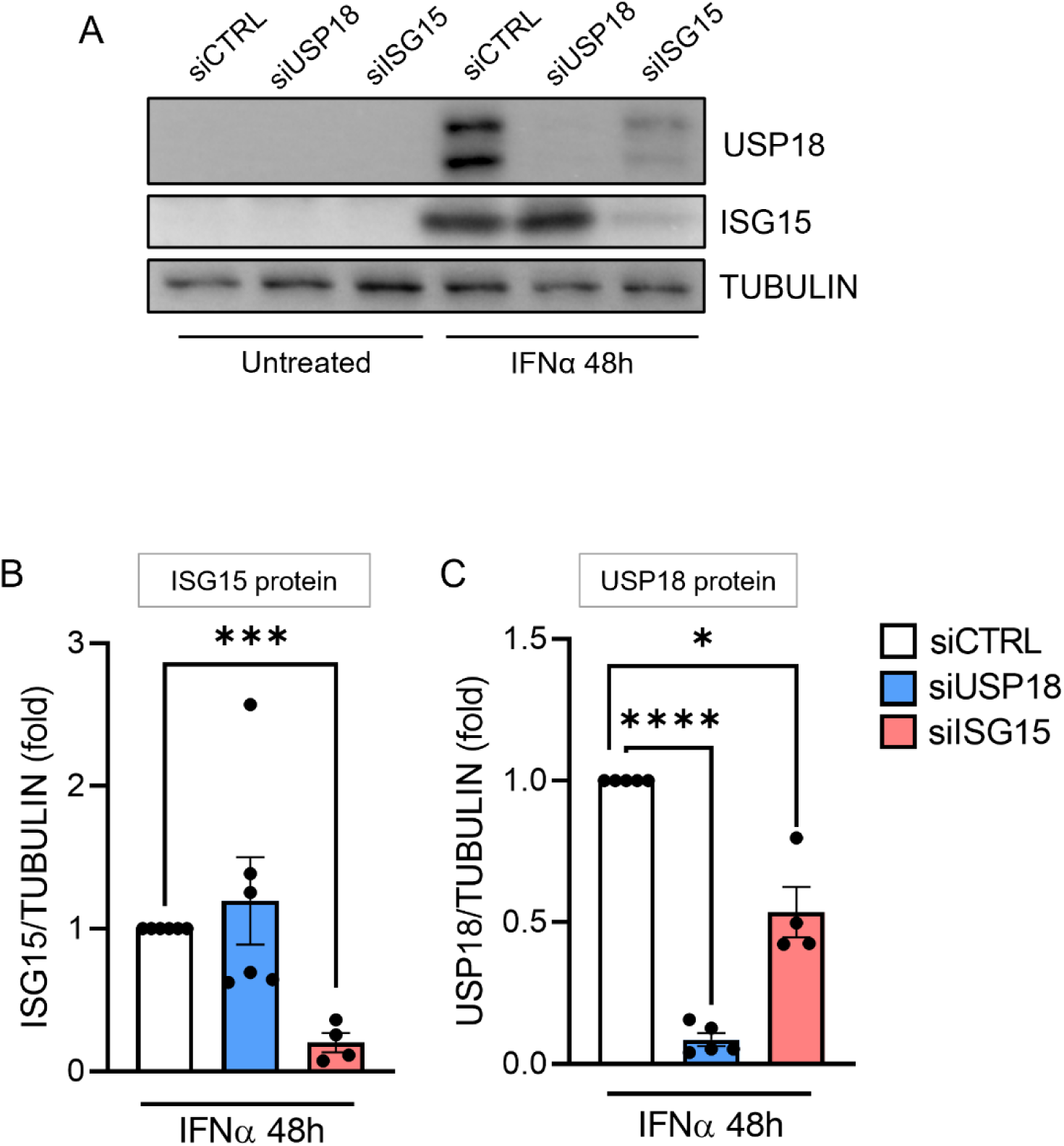
Silencing *ISG15* decreases USP18 protein expression in EndoC-βH1 cells. EndoC-βH1 cells were transfected with small interfering RNAs (siCTRL, siUSP18 or siISG15). After recovery, cells were left untreated or exposed to IFNα (2,000U/mL) for 48h. **(A)** ISG15 and USP18 protein expression were analyzed by immunoblot. **(B)** ISG15 and **(C)** USP18 bands were quantified by densitometry, normalized to TUBULIN and represented as fold compared to siCTRL IFNα-treated cells. Results are means ± SEM. Each point represents an independent experiment, i.e., independent passage. *p<0.05, ***p<0.001, ****p<0.0001 vs siCTRL, one-way ANOVA followed by Bonferroni’s post hoc test.

### *ISG15* modulates human islet cells responses to coxsackievirus B1 (CVB1) infection

We explored the role of *ISG15* in islet cells upon infection with the diabetogenic virus coxsackievirus B1 (CVB1) (52). In EndoC-βH1 cells, *ISG15* levels were not affected by CVB1 infection and the siISG15 decreased *ISG15* expression by 40% in CVB1-infected cells (not significantly, **Figure 7A**). In human islets, CVB1 infection upregulated *ISG15* mRNA and the siISG15-induced KD was about 60% (**Figure 7D**). Importantly, silencing *ISG15* enhanced the cytopathogenic effect induced by CVB1 in EndoC-βH1 cells (**Figure 7B**) and human islets (**Figure 7E**). In addition, viral titers were increased in both cell models when silenced for *ISG15* and infected with CVB1 (**Figure 7C, F**). For the *USP18* KD, no significant changes in the outcome of CVB1 infection were observed (data not shown), indicating that the antiviral roles are exclusively conveyed by *ISG15*. In mirror experiments, we next overexpressed *ISG15* in EndoC-βH1 cells to assess whether viral infection was hampered in this condition. Adeno-associated viral overexpression (OE) led to a 5-fold increase in ISG15 protein expression in untreated cells (vs the 40-fold induction by IFNα exposure, **Figure S8**). In CVB1-infected cells, *ISG15* OE levels were similar as for untreated cells (**Figure 7G, H**) and increased *ISG15* levels led to reduced cytopathogenic effect (**Figure 7I**) and viral titers (**Figure 7J**).

**Figure 7.**
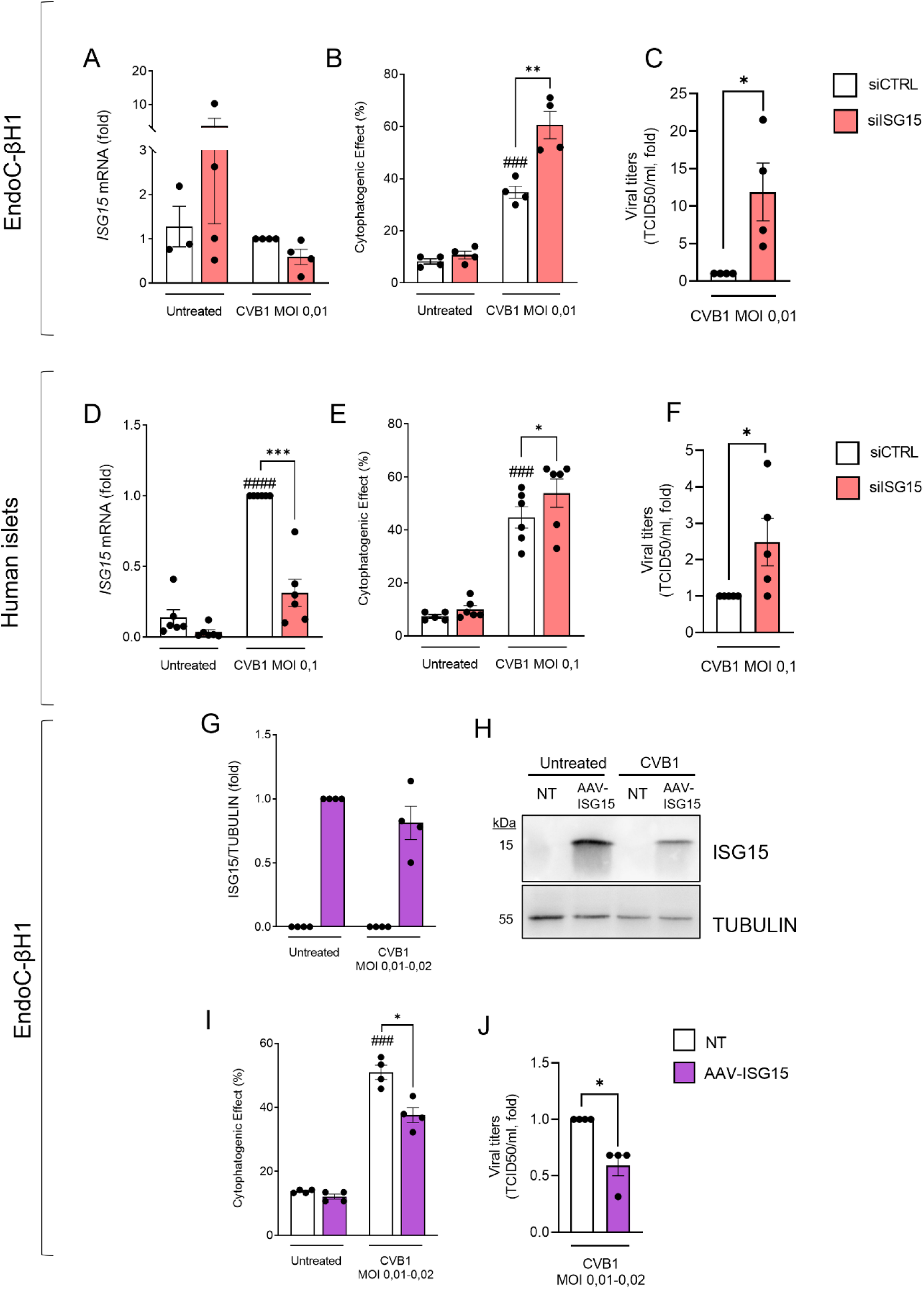
*ISG15* modulates human beta cells responses to infection by CVB1. **(A-C)** EndoC-βH1 cells and **(D-F)** human islets were transfected with small interfering RNAs (siCTRL or siISG15). After recovery, cells were left untreated or infected with CVB1 (MOI 0,01 for EndoC-βH1 cells or MOI 0,1 for human islets). **(G-J)** EndoC-βH1 cells were transduced with AAV overexpressing ISG15 (AAV-ISG15) at a MOI of 10E^5^. After recovery, cells were left untreated or infected with CVB1 (MOI 0,01-0,02). **(A, D)** *ISG15* mRNA expression was assessed by RT-qPCR, normalized to the expression of *ACTIN* and *VAPA* and represented as fold change compared to untreated cells. **(G, H)** ISG15 protein expression was analyzed by immunoblot. ISG15 bands were quantified by densitometry, normalized to TUBULIN and represented as fold compared to uninfected AAV-ISG15-treated cells. **(B, E, I)** The cytopathogenic effect was evaluated via staining with the DNA-binding dyes propidium iodide and Hoechst 33342. **(C, F, J)** Viral titers were determined in the supernatant of the CVB1 infected cells in **(B, E, I).** Results are means ± SEM. Each point represents an independent experiment, i.e., independent passage or islet donor. *p<0.05, **p<0.01, ***p<0.001 vs siCTRL, ###p<0.001, ####p<0.0001 vs untreated one-way ANOVA followed by Bonferroni’s post hoc test. Figures **C**, **F** and **J**: Unpaired t-test. CVB1: Coxsackievirus B1. MOI: multiplicity of infection. NT: non-treated, TCID: Tissue Culture Infectious Dose, AAV: Adeno-associated virus

These results confirm that *ISG15* plays an important role in the outcome of CVB1 infection in pancreatic beta cells.

## DISCUSSION

Pancreatic beta and alpha cells are both exposed to inflammation during T1D (2). However, beta cells but not alpha cells die. This difference, plus the fact that alpha cells are more resistant than beta cells to metabolic- and immune-mediated diabetogenic insults (3, 4, 61), suggests that valuable lessons for beta-cell protection can be learned from alpha cells. We recently showed that alpha cells exhibit enhanced “immune-like” gene expression compared to beta cells, not only in the context of islet inflammation but also under basal conditions, with a preponderance for expression of anti-inflammatory and anti-apoptotic genes (3, 47). The basal expression of immune genes and pathways in alpha cells, including *MDA5*, *HLA-A*, *HLA-B* and *B2M* as well as genes related to interferon signaling, in parallel to higher expression of anti-inflammatory and anti-apoptotic mRNAs, may represent an evolutionary adaptive mechanism that allows them to sense danger signals while preventing excessive immune activation, contributing to their increased survival in autoimmune diabetes. Similarly, studies have shown that bats limit excessive inflammatory cytokine expression, modulate NF-κB signalling and suppress inflammasome activation to reduce immunopathology while maintaining antiviral responses (14, 15). Bioinformatic comparisons performed herein between human islet cells and bat fibroblasts confirmed that alpha cells exhibit greater transcriptomic similarity to the antiviral responses of *Rousettus aeqyptiacus* skin fibroblasts than beta cells. Moreover, comparing alpha and beta cells’ transcriptomic data with genes associated with viral resistance and tolerance programs to infection revealed that alpha and beta cells show similar tolerance-related transcriptional profiles, but alpha cells have a higher intrinsic resistance to viral or inflammatory stress. This intrinsic resistance may make alpha cells more resistant to diabetogenic conditions. These observations point to convergent evolutionary mechanisms shared by human pancreatic alpha cells and bat skin cells, endowing alpha cells with a stronger antiviral phenotype compared to beta cells.

*ISG15* plays an important role in bats’ viral resistance (15). We re-analysed five scRNA-seq datasets and found a higher *ISG15* expression in alpha cells compared to beta cells, with a larger proportion of *ISG15*-expressing alpha cells versus beta cells. By using three different human islet-cell models, namely EndoC-βH1 cells, hiPSC-derived islet-cells and native human islets, we confirmed the higher expression of *ISG15* in alpha vs beta cells and found that *ISG15* KD under both basal and inflammatory conditions sensitized pancreatic islet cells to apoptosis and exacerbated IFNα responses, i.e., expression and secretion of the chemokine CXCL10 and expression of *HLA-ABC*. The hyperexpression of HLA-ABC in beta cells is a hallmark of T1D, promoting neoantigen presentation and immune-mediated recognition by CD8^+^ T-cells (62, 63).

Besides its roles as an IFN-I-induced gene, ISG15 controls numerous cellular pathways, including DNA damage responses (64) and intracellular trafficking (60, 65) (reviewed in (66)). The sensitization to apoptosis induced by *ISG15* KD even under basal conditions (where there is limited *ISG15* expression) suggests that *ISG15* is essential for pancreatic islet cell physiology. Our results also show that *USP18* plays a role in IFNα responses as its KD induced a similar phenotype as *ISG15* KD.

The antiviral activity of ISG15 was first observed in a screen designed to identify relevant antiviral interferon-stimulated genes during Sindbis virus infection (67). Several viruses, including influenza virus, vaccinia virus, vesicular stomatitis virus and Sendai virus have been reported to be inhibited by ISG15 *in vitro* (68). ISG15 is also essential for coxsackievirus B3 (CVB3) infection control: ISGylation in non-hematopoietic cells is required for protection from CVB3 pathology and upregulation of antiviral IFIT1/3 proteins (69). ISG15 is also reported to conjugate with the viral protease 2A, inhibiting its activity and thereby protecting mice from CVB3-induced cardiomyopathy (70). The role of ISG15 upon infection by the diabetogenic CVB1 in islet cells has not been previously explored. *ISG15* gene and protein expression was not affected by CVB1 infection in EndoC-βH1 (**Figure 7A, G, H**) but in human islets, CVB1 infection increased *ISG15* expression. This may reflect the contribution of other islet cell types to the antiviral response. In both models, *ISG15* silencing increased the cytopathogenic effect and viral production in CVB1-infected cells while its overexpression hampered the viral-induced damage in EndoC-βH1 cells. ISG15-reduces viral loads through a direct interference with viral proteins required for replication, as occurs with human papillomavirus infection (71, 72).

The canonical functions of ISG15 are executed intracellularly, but ISG15 can also be secreted to the extracellular space where it enhances the secretion of pro-inflammatory cytokines, such as IFNγ, via signalling through the leukocyte function-associated antigen (LFA-1) integrin receptor (73). This contributes to hyperinflammation during severe COVID-19 (74, 75). ISG15 exists both as a monomer and dimer, and although only the monomer can be used for intracellular ISGylation, the formation of homodimers, stabilized by disulfide bonds with a conserved cysteine (Cys78 in humans), is required for the extracellular cytokine function of ISG15 (76, 77). Interestingly, Cys78 is deleted in *Rhinolophid* and *Hipposiderid* bats, preventing ISG15 homodimerization; this decreases extracellular ISG15 cytokine-like activity while enhancing intracellular ISGylation. This may explain why bats have reduced viral-induced systemic inflammation while preserving the ability to clear the viral infection (15). ISG15 secretion from alpha cells is higher than from beta cells, and it is conceivable that secreted ISG15 by alpha cells impacts the beta cell population and contributes to local inflammation.

We observed that silencing of *ISG15* upregulates *USP18* mRNA while downregulating USP18 protein. Similar results were reported in fibroblasts from ISG15-deficient patients treated with IFNα, where *USP18* transcript levels increased but protein levels decreased (78). This may occur because intracellular free ISG15 prevents USP18 ubiquitination and degradation, promoting its stability. These findings support a post-transcriptional regulation of USP18 by ISG15.

In conclusion, by performing comparative analysis between human alpha/beta cells and bats and using different *in vitro* models of human pancreatic islet cells, *ISG15* was identified as a potential key gene in response to IFNα-mediated damage and CVB1 infection. Further comparisons of ‘super-resistance’ features present in bats and alpha cells may identify additional relevant features which may be extrapolated as therapeutic strategies to increase beta cell survival in T1D.

## ACKNOWLEDGMENTS

We thank A. Augenlicht, A. Musuaya, J. Capitaine, A. Bilheu, I. Millard, A.N. Belmahjoubi and N. Pachera (ULB Center for Diabetes Research, Université Libre de Bruxelles, Brussels, Belgium) for their excellent technical support. We sincerely appreciate the HPAP Database Consortium for providing public access to the raw scRNA-seq data of human islets. The main support to D.L.E. for this project was from grants from the Breakthrough type 1 diabetes Grant Keys: 3-SRA-2023-1379-S-B and 3-IND-2024-1549-I-X. Additional support to D.L.E. was provided by the National Institutes of Health Human Islet Research Network Consortium on beta-Cell Death & Survival from Pancreatic beta-Cell Gene Networks to Therapy [HIRN-CBDS] (grant U01 DK127786), the National Institutes of Health Grants, NIDDK, grants RO1DK126444 and RO1DK133881-01, the Win4Excellence – Gene therapy in Wallonie (GT4Health_20230920) and by a research grant from Novo Nordisk A/S.

E.M.V. was supported by the Fonds de la Recherche Scientifique (F.R.S.-FNRS, grant ID 40024397) and the Breakthrough type 1 diabetes (FY25 Fellowship ID 3-PDF-2025-1671-A-N). X.Y. was supported by the Wallonie-Bruxelles International (SUB|2020|482269; SUB|2021|518859; SUB|2022|563726) and the China Scholarship Council (201908440236). M.F.B. was supported by “Les amis des Instituts Pasteur à Bruxelles” and by the Dutch Diabetes Foundation (#2018.10.002). E.M. was supported by grant PI22/00334 from Instituto de Salud Carlos III of Spain, co-financed by the European Regional Development Fund (ERDF). A.O. was supported by the Fonds de la Recherche Scientifique (F.R.S.-FNRS, grant ID CDR 40028175) and by SFD (La Société Francophone du Diabète) (SFD-ADJ2023), France.

## DECLARATION OF INTERESTS

S.J. and A.C.T. are employees of InSphero AG, a company commercializing human islet microtissues and related services. B.Y. was a member of the management team of InSphero AG., D.L.E. is a member of the Scientific Advisory Board of InSphero AG.

E.I. is an employee of Novo Nordisk A/S. J.D.W. was an employee of Novo Nordisk, Inc.

## CONTRIBUTORS

Conceptualization: D.L.E., E.M.V. and X.Y.; Methodology: E.M.V., X.Y., M.F.B., T.H.; Investigation: E.M.V., X.Y., M.F.B., S.J., P.Z., A.R.R., J.G.O., J.M.C.J., E.I., J.D.W., A.C.T., B.Y., D.L.E.; Writing of original draft: E.M.V., X.Y., M.F.B. and D.L.E.; Funding acquisition: D.L.E; Resources: F.P., J.K-C., M.N., E.M., A.C.T., B.Y.; Supervision: D.L.E. All authors have reviewed and approved the final version of the manuscript. E.M.V. is the guarantor of the data provided. The authorship order among the 3 co-first authors was assigned based on their relative contribution to the study.

## ETHICS

Human pancreatic islets from 6 non-diabetic organ donors were isolated at the Hospital Universitari de Bellvitge - IDIBELL, Barcelona, Spain, or at the CHU Lille, France, with written consent from donors’ next-of-kin and approval of the local ethics committee. The differentiation of hiPSCs into islet-like cells was approved by the Ethics Committee of the Erasmus Hospital, Université Libre de Bruxelles, reference P2019/498.

## DATA AVAILABILITY

All original single-cell and bulk sequencing datasets used in this study are publicly available through the Human Pancreas Analysis Program (HPAP) portal (https://hpap.pmacs.upenn.edu/explore/download?matrix) or the Gene Expression Omnibus (GEO) under accession numbers: <u>GSE203384</u>, <u>GSE190726</u> and <u>GSE200083</u>. The scRNA-seq analysis workflow used in this study follows established practices and does not involve the development of new computational algorithms. All software dependencies and R package versions are reported in the Methods section. The pipeline is adapted from the publicly available HPAP workflow (https://hpap.pmacs.upenn.edu/explore/workflow/scRNAseq-analysis) and was extended to incorporate ambient RNA decontamination, Harmony-based data integration and differential expression analysis. The customized analysis scripts are stored in a private GitHub repository (https://github.com/xiaoyanyi666/HPAPplus-scRNA-seq-pipeline-with-decontamination-and-DEG-analysis) and are available from the authors upon reasonable request.

