## Supplemental Figures for "From bats to humans: uncovering *ISG15* as a new resistance factor in type 1 diabetes"

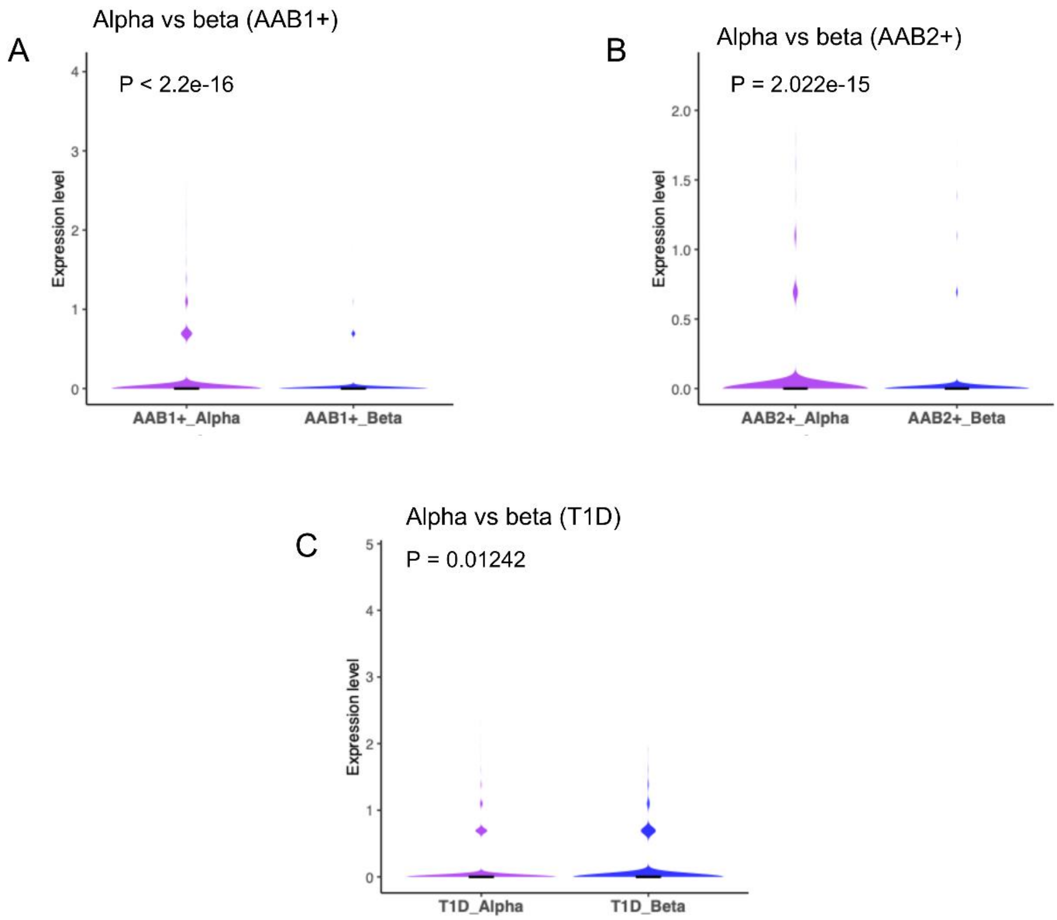


**Figure S1. *ISG15* expression in alpha cells and beta cells in AAB+ and T1D donors.** Violin plots with embedded boxplots showing single-cell RNA-seq expression in primary islets from HPAP AAB1+ donors **(A)**, AAB2+ donors **(B)** and T1D donors **(C)**. Adjusted P-values (Wilcoxon rank-sum test with Benjamini-Hochberg correction) for panels **(A-C)** are indicated. AAB+: autoantibody positive (1 or more). HPAP: Human Pancreas Analysis Program (https://hpap.pmacs.upenn.edu/), T1D: Type 1 diabetes.

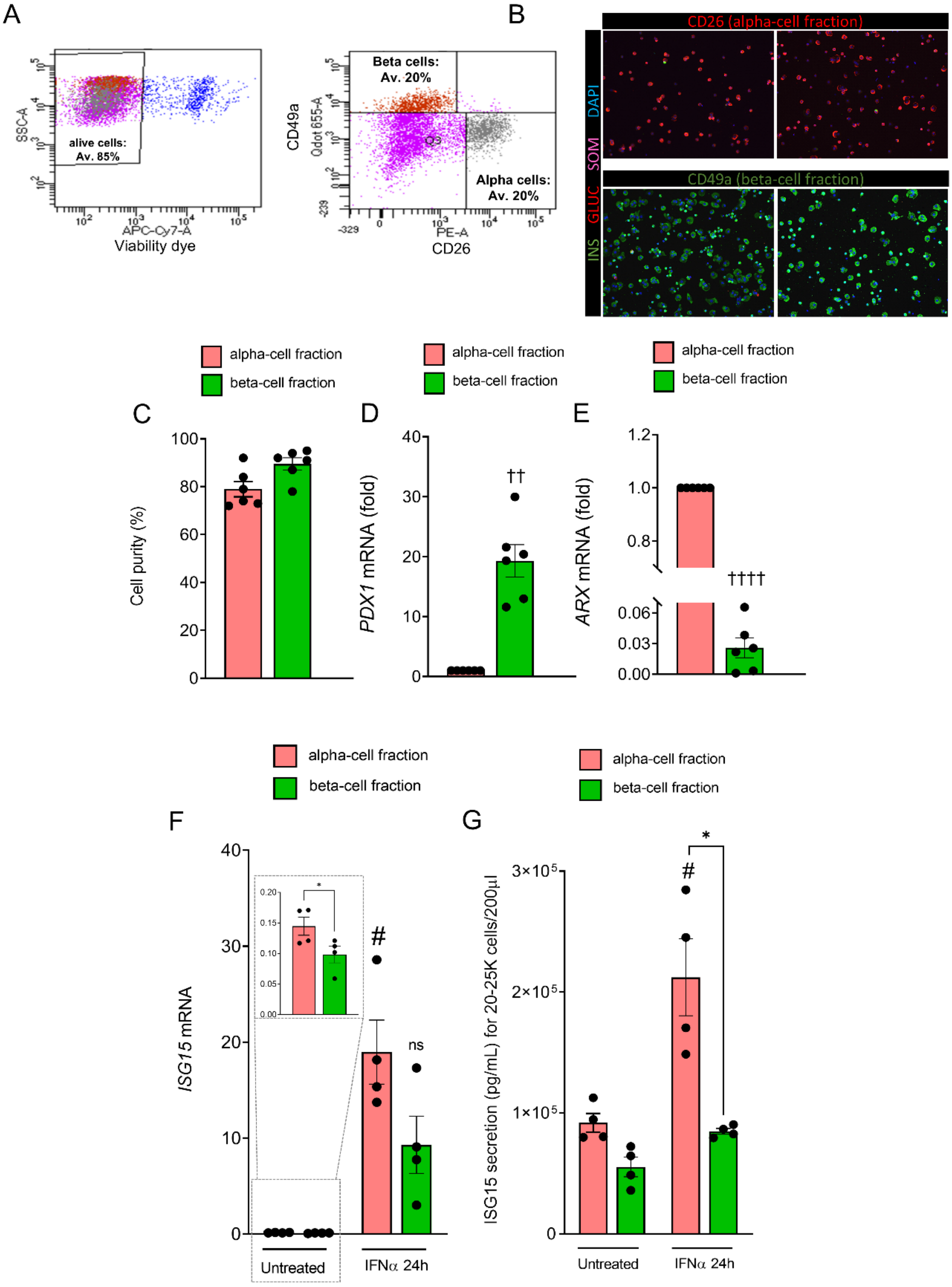


**Figure S2. Validation of *ISG15* increased expression and secretion in hiPSC-derived and purified alpha cells compared to beta cells. (A)** Islet-like aggregates derived from the hiPSC line 1023A were further differentiated by an extra 4 weeks of culture as described [1]. Alive cells were then purified by FACS using a viability dye and enrichment of alpha/beta cells was done using the antibodies CD26-PE for alpha cells and CD49a-FITC for beta cells. **(B)** Representative immunofluorescence images post-sorting co-stained for insulin (INS, green), glucacon (GCG, red) and somatostatin (SOM, pink). Nuclei were stained with DAPI (blue). **(C)** Quantifications of INS+ (beta) and GCG+ (alpha) cells from the immunofluorescence pictures were performed to determine the purity of the post-sorting populations. **(D)** *PDX1* and **(E)** *ARX* mRNA expression was assessed by RT-qPCR in purified alpha and beta cells, normalized to the geometric mean of *ACTIN* and *VAPA* and presented as fold change compared with alpha cells. **(F, G)** Purified alpha and beta cells were exposed or not to IFNα (2000 U/ml) for 24h. **(F)** *ISG15* mRNA expression was assessed by RT-qPCR and normalized to the geometric mean of *ACTIN* and *VAPA*. **(G)** ISG15 secretion to the supernatant was determined by ELISA. Results are means ± SEM. Each point represents an independent experiment, i.e., independent differentiation. *$p<0.05$ vs alpha cells exposed or not to IFNα, #$p<0.05$ vs untreated one-way ANOVA followed by Bonferroni's post hoc test. ††$p<0.01$, ††††$p<0.0001$ vs alpha cells Student t-test.

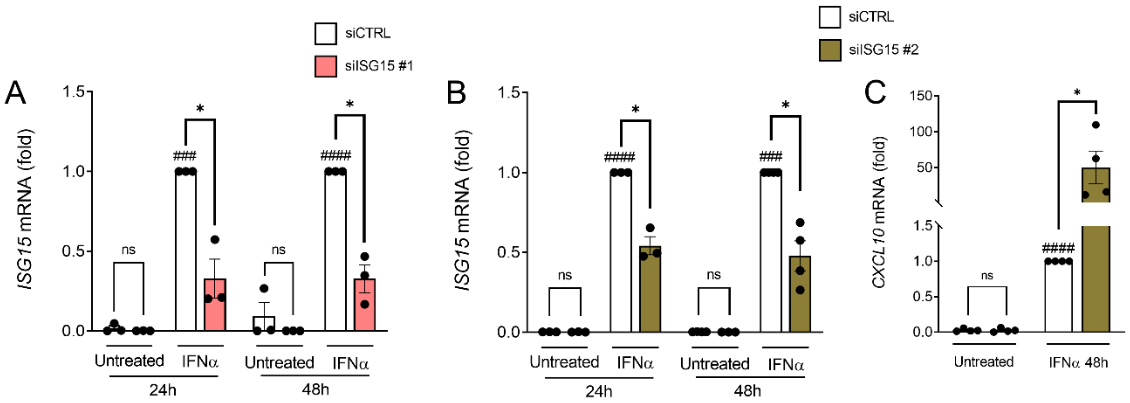


**Figure S3. Validation of *ISG15* silencing in EndoC-βH1 cells.** EndoC-βH1 cells were transfected with small interfering RNAs (siCTRL, siISG15 #1 or siISG15 #2). After 24h of recovery, cells were left untreated or exposed to IFNα (2,000U/ml) for 24h or 48h. **(A, B)** *ISG15* and **(C)** *CXCL10* mRNA expression (after siISG15 #2) was assessed by RT-qPCR, normalized to the geometric mean of *ACTIN* and *VAPA* and presented as fold change compared with siCTRL-IFNα-treated cells. SiISG15 #1 was used in all subsequent experiments (See ***Figure 5A*** for *CXCL10* mRNA expression after siISG15 #1). Results are means ± SEM. Each point represents an independent experiment, i.e., independent passage. *p<0.05 vs siCTRL, ###p<0.001, ####p<0.0001 vs untreated one-way ANOVA followed by Bonferroni's post hoc test.

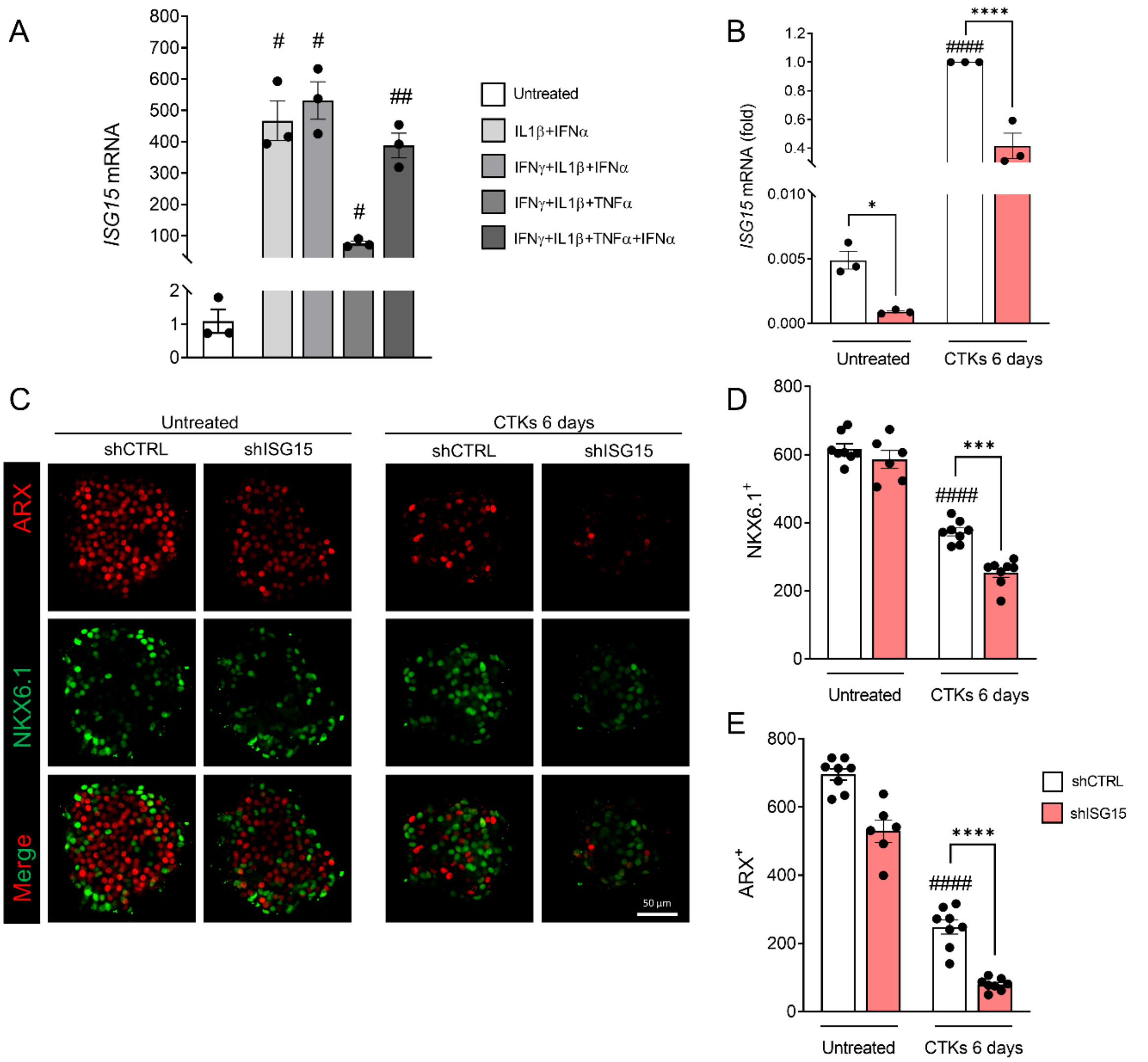


**Figure S4. Silencing *ISG15* sensitizes human islet microtissues (hIsMTs) to cytokines. (A)** hIsMTs were left untreated or exposed to different combinations of IFNγ (25ng/mL) +IL1β (5ng/mL) +TNFα (25ng/mL) +IFNα (10ng/mL) as detailed in the figure, for 6 days. **(A, B)** *ISG15* mRNA expression was assessed by RT-qPCR, normalized to the geometric mean of *ACTIN* and *VAPA*. **(B-E)** hIsMTs were transfected with short-hairpin RNAs (shCTRL or shISG15). After recovery, cells were left untreated or exposed to IFNγ (25ng/mL) +IL1β (5ng/mL) +TNFα (25ng/mL) +IFNα (10ng/mL) for 6 days. **(C)** Representative 3D immunofluorescence pictures of hIsMTs silenced for *ISG15* and exposed to cytokines stained for NKX6.1 (green) and ARX (red). **(D, E)** Quantifications of **(D)** beta- and **(E)** alpha-cell fractions were performed using customized CellPathfinder pipelines. Results are means ± SEM. Each point represents a technical replicate from a single donor. ***p<0.001, ****p<0.0001 vs shCTRL, #p<0.05, ##p<0.01, ####p<0.0001 vs untreated one-way ANOVA followed by Bonferroni's post hoc test. hIsMTs: Human islet microtissues; CTKs: cytokines.

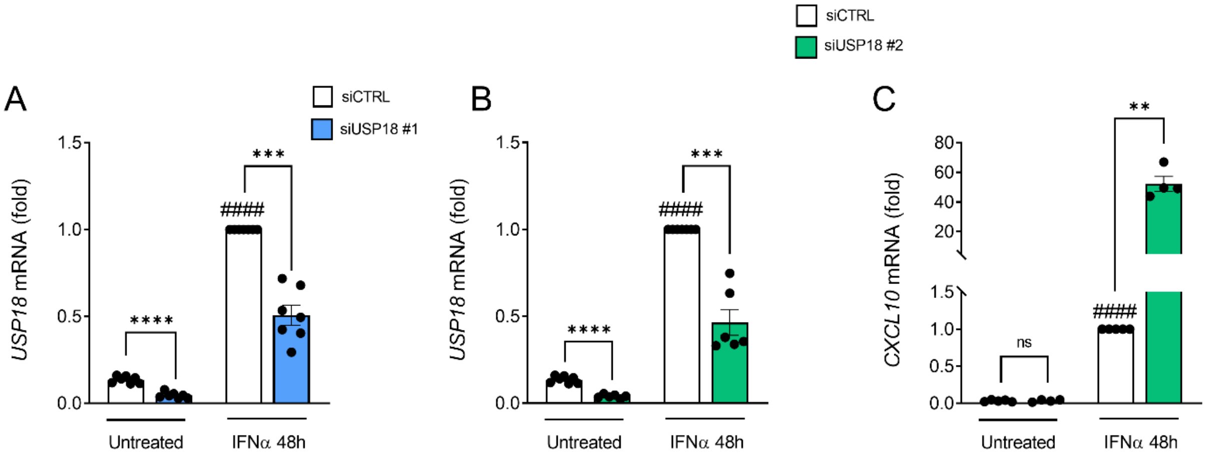


**Figure S5. Validation of *USP18* silencing in EndoC-βH1 cells.** EndoC-βH1 cells were transfected with small interfering RNAs (siCTRL, siUSP18 #1 or siUSP18 #2). After 24h of recovery, cells were left untreated or exposed to IFNα (2,000U/ml) for 48h. **(A, B)** *USP18* and **(C)** *CXCL10* mRNA expression (after siUSP18 #2) was assessed by RT-qPCR, normalized to the geometric mean of *ACTIN* and *VAPA* and presented as fold change compared with siCTRL IFNα-treated cells. SiUSP18 #1 was used in all subsequent experiments. (See ***Figure S6D*** for *CXCL10* mRNA expression after siUSP18 #1). Results are means ± SEM. Each point represents an independent experiment, i.e., independent passage. $**p<0.01$, $***p<0.001$ vs siCTRL, ####$p<0.0001$ vs untreated one-way ANOVA followed by Bonferroni's post hoc test.

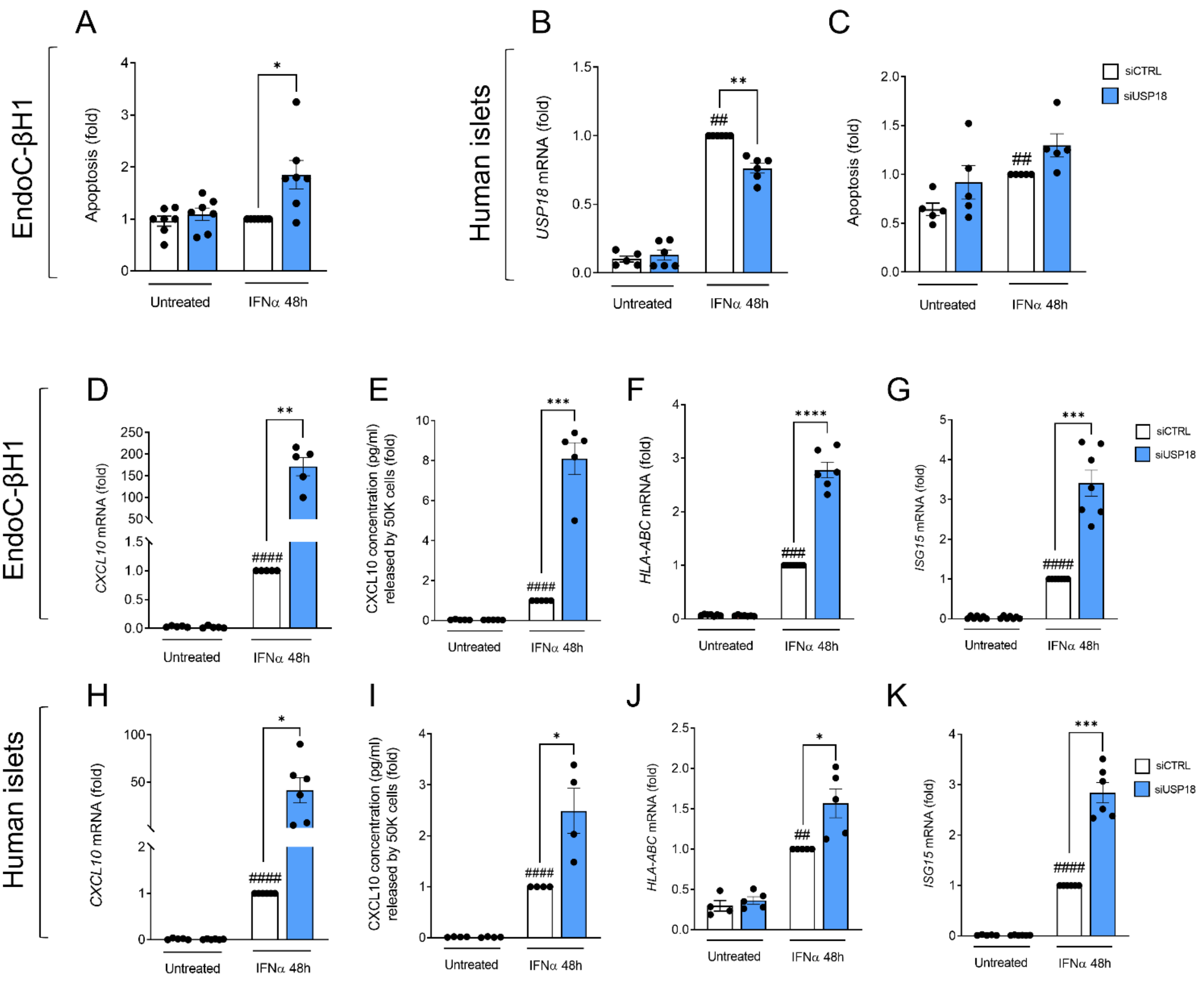


**Figure S6. Silencing *USP18* upregulates IFNα-induced downstream events. (A, D-G)** EndoC-βH1 cells and **(B, C, H-K)** human islets were transfected with small interfering RNAs (siCTRL or siUSP18). After recovery, cells were left untreated or exposed to IFNα (2,000U/mL) for 48h. **(A, C)** Apoptotic cells were counted with the DNA-binding dyes propidium iodide and Hoechst 33342. **(B)** *USP18* **(D, H)** *CXCL10*, **(F, J)** *HLA-ABC* and **(G, K)** *ISG15* mRNA expression was assessed by RT-qPCR, normalized to the expression of *ACTIN* and *VAPA* and represented as fold change compared to siCTRL IFNα-treated cells. **(E, I)** CXCL10 secretion in the supernatant was measured by ELISA. 50,000 cells were used per condition. Results are means ± SEM. Each point represents an independent experiment, i.e., independent passage or islet donor. *p<0.05, **p<0.01, ***p<0.001, ****p<0.0001 vs siCTRL, ##p<0.01, ###p<0.001, ####p<0.0001 vs untreated one-way ANOVA followed by Bonferroni's post hoc test.

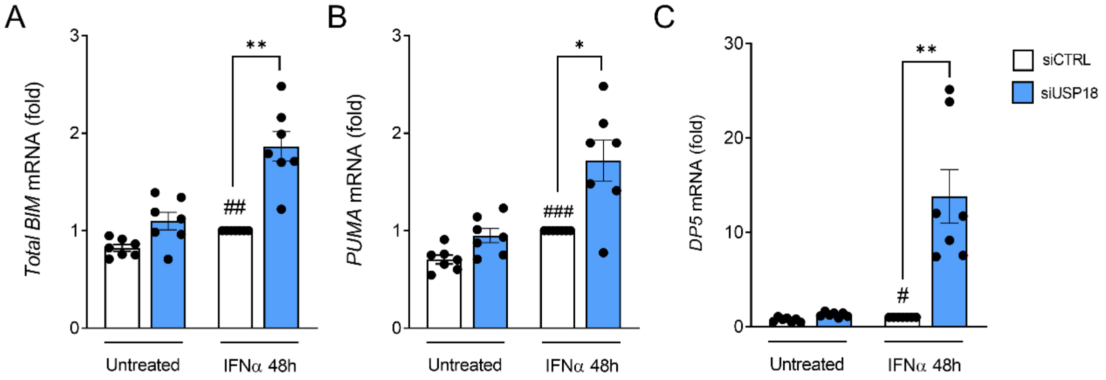


**Figure S7. *USP18* modulates the intrinsic apoptosis pathway.** EndoC-βH1 cells were transfected with small interfering RNAs (siCTRL or siUSP18). After recovery, cells were left untreated or exposed for 48h to IFNα (2,000U/mL). **(A)** *Total BIM,* **(B)** *PUMA* and **(C)** *DP5* mRNA expression was assessed by RT-qPCR, normalized to the expression of *ACTIN* and *VAPA* and represented as fold change compared to siCTRL IFNα-treated cells. Results are means ± SEM. Each point represents an independent experiment, i.e., independent passage. *p<0.05, **p<0.01vs siCTRL, #p<0.05, ##p<0.01, ###p<0.001 vs untreated, one-way ANOVA followed by Bonferroni's post hoc test.

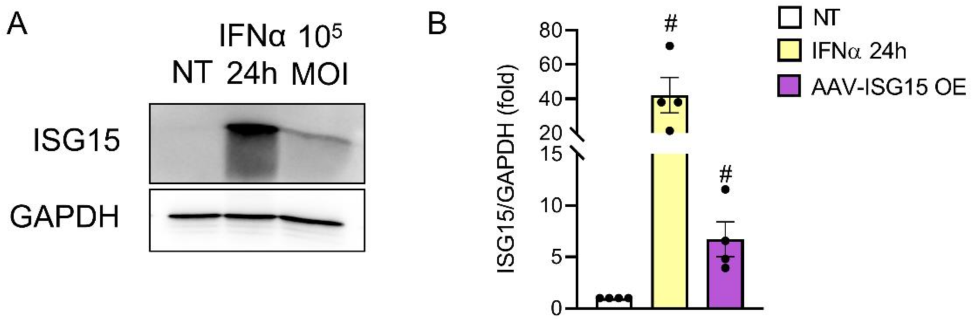


**Figure S8. Validation of ISG15 overexpression in EndoC-βH1 cells.** EndoC-βH1 cells were transduced with AAV overexpressing ISG15 (AAV-ISG15 OE) at a MOI of $10E^5$. In parallel, uninfected cells were exposed or not to IFNα (2,000U/mL) for 24h. **(A)** ISG15 and GAPDH protein expression were analyzed by immunoblot. **(B)** ISG15 bands were quantified by densitometry, normalized to GAPDH and represented as fold compared to uninfected and untreated cells. Results are means ± SEM. Each point represents an independent experiment, i.e., independent passage. #p<0.05 vs NT, one-way ANOVA followed by Bonferroni's post hoc test. AAV: Adeno-associated virus, MOI: multiplicity of infection, NT: non-treated, OE: overexpression.
