## Supplemental Table S4-S5 for "From bats to humans: uncovering *ISG15* as a new resistance factor in type 1 diabetes"

| Data set | ID | Description | Data type | Tissue | Samples | Age | Source | Year |
|---|---|---|---|---|---|---|---|---|
| 1 | HPAP-young (non-diabetic) | HPAP-young (non-diabetic) | scRNA-seq | Pancreatic islets | 15 (7F/8M) | 22.7 ±12.8 | HPAP [1] | 2025; 2022 |
| 2 | HPAP-older (non-diabetic) | HPAP-older (non-diabetic) | scRNA-seq | Pancreatic islets | 13 (7F/6M) | 42.5 ±10.7 | HPAP [1] | 2025; 2022 |
| 3 | HPAP-AAB1+ | HPAP-AAB1+ | scRNA-seq | Pancreatic islets | 9 (3F/6M) | 43.1 ±10.5 | HPAP [1] | 2025; 2022 |
| 4 | HPAP-AAB2+ | HPAP-AAB2+ | scRNA-seq | Pancreatic islets | 2 (2M) | 48±15.6 | HPAP [1] | 2025; 2022 |
| 5 | HPAP-young (T1D control) | HPAP-young (T1D control) | scRNA-seq | Pancreatic islets | 15 (7F/8M) | 22.7 ±12.8 | HPAP [1] | 2025; 2022 |
| 6 | Szymczak et al. | iPSC stage 7 | scRNA-seq | hiPSC-derived islet like cells | 5 | / | GSE203384 [2] | 2022 |
| 7 | Chandra et al. | iPSC stage 6 | scRNA-seq | hiPSC-derived islet like cells | 1 | / | GSE190726 [3] | 2022 |
| 8 | Sintov et al. (*in vivo*) | hESC *in vivo* | scRNA-seq | hESC-derived islet like cells | 5 | / | GSE200083 [4] | 2022 |
| 9 | HPAP-basal | HPAP-basal | Bulk RNA-seq | FACS-sorted alpha-and beta cells | 12 (4F/8M) | 35.3 ±10.8 | HPAP [1] | 2024 |

**Table S4. Overview of the RNA-seq metadata used in the present study.** RNA-seq data for basal alpha- and beta cells were obtained from the Human Pancreas Analysis Program (HPAP, https://hpap.pmacs.upenn.edu/) and the Gene Expression Omnibus (GEO) portal. Age is presented as mean ± SD (when available for the donors). F: female; M: male; hiPSC: human induced pluripotent stem cells; hESC: human embryonic stem cells.

| Islet prep. | 1 | 2 | 3 | 4 | 5 | 6 |
|---|---|---|---|---|---|---|
| MANDATORY INFORMATION | | | | | | |
| Unique identifier | 215.25 | 216.25 | HI341-A | HI345 | HI351 | HI359-A |
| Donor age (years) | 77 | 53 | 54 | 61 | 57 | 56 |
| Donor sex (M/F) | M | M | F | F | M | M |
| Donor BMI (kg/m$^2$) | 22.04 | 25.47 | 34.1 | 22 | 27.2 | 22 |
| Donor $HbA_{1c}$ or other measure of blood glucose control | Hba1c: 5.4% | Hba1c: 5.4% | Hba1c: 5.8% | Hba1c: 6.2% | Hba1c: 5.7% | Hba1c: 5.9% |
| Origin of islets | L'Hospitalet de Llobregat, Barcelona, Spain | L'Hospitalet de Llobregat, Barcelona, Spain | Diabetes, University of Lille, Lille, France | Diabetes, University of Lille, Lille, France | Diabetes, University of Lille, Lille, France | Diabetes, University of Lille, Lille, France |
| Islet isolation centre | Laboratori de la Unitat de Trasplantament d'Illots Pancreàtics (UTIP) | Laboratori de la Unitat de Trasplantament d'Illots Pancreàtics (UTIP) | European Genomic Institute for Diabetes, Translational Research for Diabetes | European Genomic Institute for Diabetes, Translational Research for Diabetes | European Genomic Institute for Diabetes, Translational Research for Diabetes | European Genomic Institute for Diabetes, Translational Research for Diabetes |

| | | | | | | |
|---|---|---|---|---|---|---|
| **Donor history of diabetes? Please select yes/no from drop down list** | No | No | No | No | No | No |
| **RECOMMENDED INFORMATION** | | | | | | |
| **Donor cause of death** | Anoxia after Cardiac arrest | Cardiac arrest | Stroke | Stroke | Stroke | Stroke |
| **Estimated purity (%)** | 40 | 47 | 70 | 68 | 53 | 47 |
| **Total culture time** | 8 days | 8 days | 8 days | 8 days | 8 days | 8 days |

**Table S5.** Checklist for reporting human islet preparations used in research adapted from [5], related to **Figures 4, 5, 7, S6**. Estimated purity is based on staining for insulin as described in Methods.
